# Dynamic microfluidic single-cell screening identifies pheno-tuning compounds to potentiate tuberculosis therapy

**DOI:** 10.1101/2023.03.31.535085

**Authors:** Maxime Mistretta, Mena Cimino, Pascal Campagne, Stevenn Volant, Etienne Kornobis, Olivier Hebert, Christophe Rochais, Patrick Dallemagne, Cédric Lecoutey, Camille Tisnerat, Alban Lepailleur, Yann Ayotte, Steven R. LaPlante, Nicolas Gangneux, Monika Záhorszká, Jana Korduláková, Sophie Vichier-Guerre, Frédéric Bonhomme, Laura Pokorny, Marvin Albert, Jean-Yves Tinevez, Giulia Manina

## Abstract

Drug-recalcitrant infections are a leading global-health concern. Bacterial cells benefit from phenotypic variation, which can suggest effective anti-microbial strategies. However, probing phenotypic variation entails spatiotemporal analysis of individual cells that is technically challenging, and hard to integrate into drug discovery. To address this, we developed a flow-controlled multi-condition microfluidic platform suitable for imaging two-dimensional growth of bacterial cells, compressed inside separate microchambers by a soft hydro-pneumatic membrane. With this platform, we implemented a dynamic single-cell screening for compounds that induce a phenotypic change while decreasing cell-to-cell variation, aiming to undermine the bacterial population, making it more vulnerable to other drugs. We first applied this strategy to mycobacteria, as tuberculosis poses a major public-health threat. Our top hit impairs *Mycobacterium tuberculosis* via a peculiar mode of action and enhances other anti-tubercular drugs. This work proves that pheno-tuning compounds represent a successful approach to tackle pathogens that are increasingly difficult to treat.

## Introduction

The advent of antimicrobials revolutionized medical practice, saving countless lives, but bacterial pathogens were not slow to develop drug-evasion mechanisms^1^. The common factor among them is the unique ability of microbes to diversify, increasing their chances of survival^2^. Genetic changes occur rarely and confer a stable advantage to the whole population. Phenotypic changes occur regularly and confer a transient advantage to a subset of individuals, but can have a positive effect on whole-population fitness^3^; they are multifactorial and can possibly evolve into stable heritable changes^4^. Overall, the marked diversification potential of microbial pathogens, in addition to physicochemical barriers to drug penetration and inefficient host clearance, largely contributes to persistent and recurrent infections and is conducive to treatment failure^5, 6^. As a result, nearly a century after the golden era, the world is running into an alarming post-antibiotic era, which is jeopardizing global-health stability^7^. This emphasizes the pressing need to tackle bacterial pathogens with original solutions, which reckon with the inherent and environment-driven phenotypic variation of clonal microbial populations^8^.

A leading example of public-health threat is *Mycobacterium tuberculosis*, which can persist for months up to years in the host despite protracted therapy and is responsible for about a quarter of the antimicrobial resistance emergency^9^. Although some new and repurposed drugs have been introduced in recent years^10, 11^, tuberculosis treatment remains a slow and grueling process that requires improvement. Shortening treatment duration and decreasing relapse rates entail targeting drug-tolerant and persistent subsets, by either preventing their formation or hindering their survival mechanisms^6, 8, 12–14^. Nevertheless, the underlying limitation of conventional drug discovery approaches, whether target- or cell-based, lies in the fact that they rely on reductionist assessments, with little-to-no resolution on minor cellular subsets, responsible for drug evasion^7, 9, 15^. Hence, selection of more effective drug candidates should rely on original decision criteria, grounded in the quantification of single-cell dynamics, and aimed at leveraging phenotypic variation towards enhanced therapeutics.

Diverse instances of phenotypic variation related to growth, division, and gene expression, as well as their association with phenotypic drug tolerance, were reported in mycobacteria^16^. A relevant case of cell-to-cell variation concerns genotoxic stress, which can occur due to intrinsic replication errors or exogenous aggression, such as immune effectors and drugs^6^. Using a fluorescent reporter of RecA as a proxy for DNA damage response, we found that more than half of clonal mycobacterial cells, grown in stress-free conditions, experiences transient DNA damage events, measured as single-cell pulses of fluorescence, which largely resolve spontaneously. In addition, pulsing cells are more likely to die upon treatment with DNA-targeting drugs, as opposed to non-pulsing cells that survive more^17^. Based on these findings, we hypothesized that manipulating RecA phenotypic variation might prevent drug-evasion mechanisms and weaken the mycobacterial population. To this aim, we sought to carry out a microfluidic dynamic single-cell screening for pheno-tuning compounds (PTC), which we refer to as µDeSCRiPTor, aiming to increase RecA expression levels and decrease RecA cell-to-cell variation, to make clonal mycobacterial cells more homogeneously susceptible to standard treatment. However, the technology required to implement this approach did not exist.

Biocompatible microfluidic devices in conjunction with time-resolved microscopy are instrumental to capture the behavior of live cells over space and time under tight environmental control, but they are complex to scale up for high-throughput applications^18^. Although digital microfluidics allows higher-throughput applications than continuous-flow microfluidics, encapsulation of small bacterial cells within hydrolipid droplets poses major limitations to both spatiotemporal resolution and environmental flexibility^19^. To overcome these drawbacks, we present here a customized multi-condition microfluidic platform, stemming from our scalable microfluidic cell-culture chamber^20^, which operates on gentle hydro-pneumatic trapping of cells and allows two-dimensional growth. In short, we combined thirty-two pairs of microchambers that not only share a common microfluidic network but are also supplied via independent reservoirs, to be able to inject a different solution into each microchamber. With this platform, we could carry out the first proof of concept of µDeSCRiPTor. By testing fewer than a hundred compounds, we were able to identify four main hits that meet our criteria for a PTC compound. Following in-depth characterization of our best hit, we found that it acts through a dual mechanism, which impairs both cell envelope and DNA, with marked increase in oxidative stress. This original drug-discovery model enabled us to identify a compound that not only undermines the mycobacterial cell per se but also enhances existing drugs, holding promise for therapeutic development against tuberculosis and other infectious diseases.

## Results

### Creation of a multi-condition microfluidic platform for long-term imaging of single cells in 2D

To implement the µDeSCRiPTor strategy, we first had to construct a multi-condition microfluidic platform compatible with bacterial growth (Fig. 1a), starting from a scalable module we developed earlier^20^. This module is composed of two layers of polymethyl siloxane (PDMS) patterned with a 300µm-wide channel crossing a 1000µm-wide circular chamber. The two layers are bonded perpendicularly to each other and in turn to a glass coverslip, creating two functional compartments: an upper control layer, where water is perfused; and a lower flow layer, where bacterial suspension and culture medium are perfused (Extended Data Figs. 1a and b). The upper section of the flow layer constitutes the third functional element of the module, namely, a 20µm-high flexible membrane on top of the circular chamber. By increasing pressure in the control layer, the membrane pneumatically lowers, forming a contact zone with the underlying coverslip, where bacteria are confined and forced to replicate in two dimensions (Extended Data Fig. 1c).

**Figure 1.**
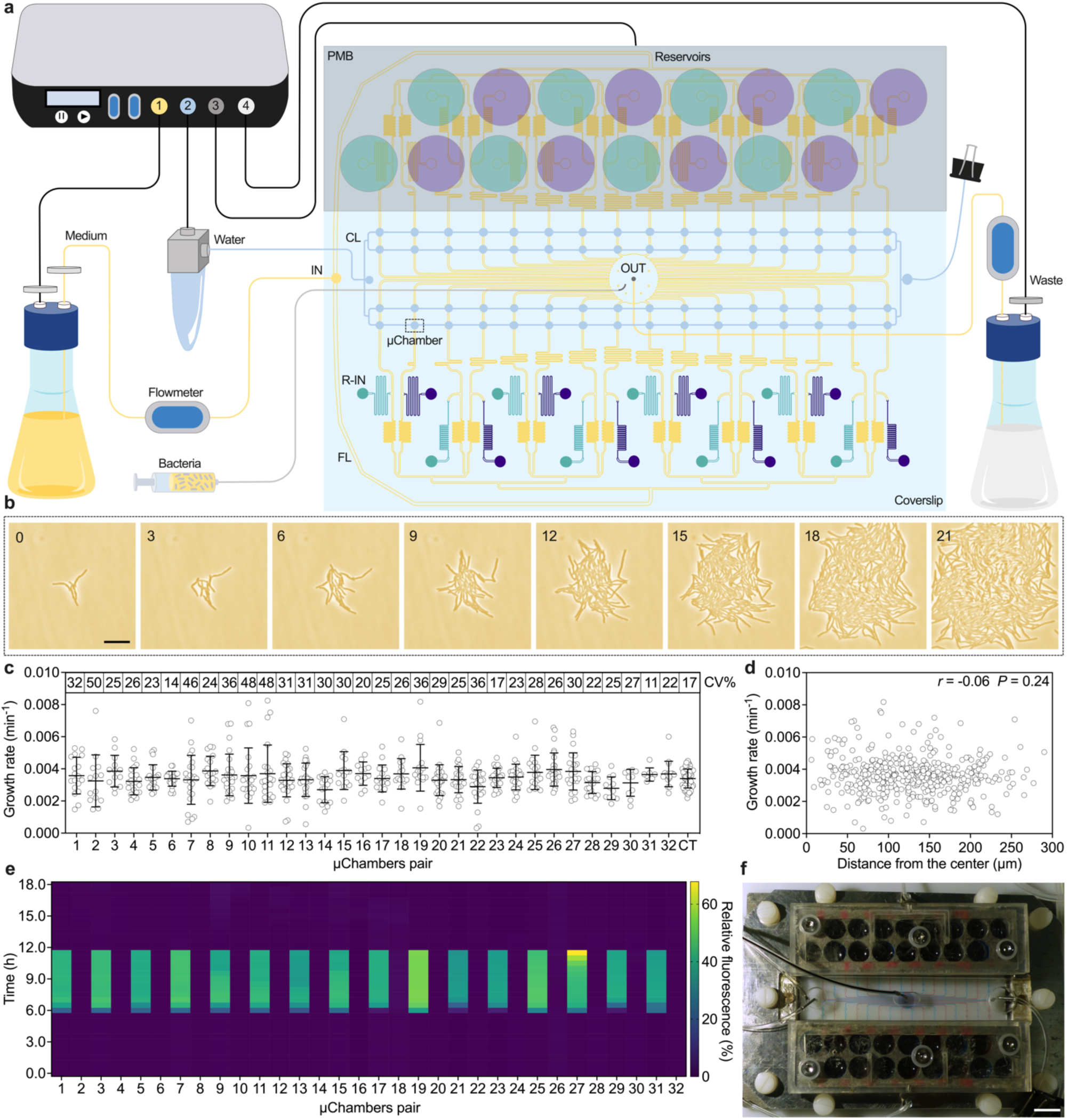
Operation and characterization of the 32-condition platform. **a**, Schematic of the 32-condition microfluidic device connected to a 4-channel software-driven flow controller. The first channel actuates the injection of medium into the flow layer (FL, yellow) by the inlet port (IN). The microfluidic network branches to 32 pairs of circular microchambers (1-mm diameter), which share the same outlet port (OUT). The OUT is connected to a waste receptacle, controlled by the fourth channel. To prevent contaminations, 0.2-µm filters (gray cylinders) are placed at the inlet and outlet ports of the medium bottle and at the waste outlet port. The OUT is also used to load the bacterial suspension. The flow of medium is regulated upstream and downstream of the device by means of two software-driven flowmeters. The FL is bound to the underlying coverslip, enabling inverted microscopy. The second channel actuates water into the control layer (CL, light blue), which is closed at one end with a binder clip. The third channel actuates the injection of different solutions from the reservoirs (larger green and purple circles) of a perforated metal block (PMB, gray rectangle). The PMB is superimposed on the reservoir inlets (R-IN, smaller green and purple circles). **b**, Representative phase-contrast image series of *M. smegmatis* growing inside one microchamber of the device (dashed square). Cells were imaged at 20-min intervals and numbers represent hours. Scale bar, 5 μm. **c**, Microcolony growth rates of *M. smegmatis* growing inside the different microchambers, compared to our control (CT) device^17^. Error bars represent mean ± SD (*n* = 688, from two independent experiments). No significant difference was found by two-way ANOVA followed by Tukey’s multiple comparisons test. Numbers in the insets indicate the percent coefficient of variation per microchamber. **d**, Spearman correlation between grow rate and position of microcolonies within the microchamber (*n* = 453, from two independent experiments). **e**, Measurement of fluorescence between each microchamber pair perfused with non-fluorescent medium for 18 hours, including a 6-hour pulse of fluorescent and non-fluorescent solutions from alternate reservoirs. Values are relative to the maximum intensity of a 100-µM concentrated FITC solution perfused into the whole network (*n* = 2). **f**, Picture of the 32-condition platform assembly, upon alternate injection of red and blue dyes from the reservoirs. Scale bar, 10 mm.

This microfluidic module, from now on referred to as microchamber, is scalable and can be multiplied and combined into diverse configurations. Here, aiming to develop a multi-condition device suitable for live screening applications, we created a flow layer made of an array of 16 x 2 pairs of microchambers overlaid by a control layer, whose microchannels are arranged perpendicular to the microchannels of the flow layer (Fig. 1a; Extended Data Fig. 1d and Source Data 1). The microchambers are fed from a single inlet port through a unique microfluidic network, which separates into 16 individual branches per side. Each branch alternates straight microchannels and microserpentines of diverse lengths, to ensure that the stream flows into the different microchambers at the same time. Each branch is also independently connected to a second separate inlet path, which in turn is fed from an individual reservoir. All microchambers are connected to an output channel that flows into a common outlet port, toward a sealed waste container. Our 32-condition device is regulated from a software-driven flow controller, via four channels (Fig. 1a). The first channel actuates the flow layer by applying a pressure of 100 mbar to a bottle of medium. The second channel actuates the control layer, which is clamped at one end, by applying a pressure of 30 mbar to a water supply, resulting in lowering the membrane on the underlying coverslip (Extended Data Fig. 1c). The third channel is connected to two arrays of reservoirs and is set at 30 mbar. The fourth channel is connected to the waste container and is set at 20 mbar. The flow rate is constantly being measured by two flowmeters installed upstream and downstream of the flow layer, and adjusted by ±15 mbar, aiming to obtain a targeted flow rate in the flow layer of 160 µL/h at the inlet and of 150 µL/h at the outlet, to prevent unwanted injection from the reservoirs.

To validate our platform, a single-cell suspension of the tuberculosis model *Mycobacterium smegmatis* was manually loaded from the outlet port, which was then connected to the waste container. Following 10-min incubation, both the control layer and the flow layer were actuated to block and feed bacterial cells, respectively. The device was mounted on the microscope stage, using a metallic and acrylic support system, held together by screws, and having a series of openings to allow insertion of other components from the top and movement of the objective from the bottom (Extended Data Figs. 1e and f). We monitored single-cell growth dynamics by inverted phase-contrast microscopy (Fig. 1b and Supplementary Video 1) and found no significant difference in microcolony growth rate across the different microchambers (Fig. 1c), regardless of the position within the PDMS-glass contact area (Fig. 1d). However, the coefficient of variation of growth rate was higher in this multi-condition device, compared with our single-condition reference device^17^, most likely due to the manual microfabrication process, resulting in intra-chamber variations of membrane adhesion and pressure fluctuations.

Finally, we integrated the fourth structural component, namely, the reservoirs, which are made of two perforated metal blocks placed on top of each side of the device (Fig. 1a and Extended Data Fig. 1g) and controlled by air injection via two sealed lids (Extended Data Fig. 1h). The 32 reservoirs, 16 holes per metal block, are interconnected to 32 inlet ports, flowing into each branch of the flow layer toward a single pair of microchambers. To check the operation of the injection system, we alternately loaded the reservoirs with fluorescent and non-fluorescent medium. We actuated the reservoirs by decreasing the pressure to 60 mbar in the first channel of the flow controller, and by increasing the pressure of the third channel from 30 to 100 mbar, resulting in simultaneous injection of the content of the 32 separate reservoirs into the microchambers. Injection was stopped by restoring the pressure to 100 mbar in the first channel and to 30 mbar in the third channel. We imaged a dynamic pattern of injection and were able to confirm the absence of cross contamination between different microchambers up to the outlet port, and the synchrony of injection and washing stages (Figs. 1e and f; Extended Data Figs. 1i−k). We also calculated that the solutions stored in the reservoirs undergo about 2.5-fold dilution during the injection phase, due to the presence of a low but steady flow of medium from the main inlet port during injection from the reservoirs.

In conclusion, we constructed and characterized a multi-layer microfluidic platform, suitable for long-term imaging of single bacterial cell growth under 32 independent conditions.

### Dynamic single-cell screening for pheno-tuning compounds (µDeSCRiPTor)

We reported that RecA, a key enzyme for DNA damage repair in the SOS response^21^, is unevenly expressed in clonal mycobacterial cells in the absence of stress, as transient pulses occurring in more than half of the population^17^. RecA-pulsing cells are also more likely to die upon treatment with DNA-damaging agents, such as fluoroquinolones and mitomycin C (MIT). Based on these findings, we hypothesized that decreasing cell-to-cell variation, by exogenous induction of RecA, could sensitize mycobacterial cells to treatment. To this aim, we leveraged our 32-condition platform to carry out an original time-resolved single-cell screening for PTC, i.e., compounds that fine-tune pre-existing phenotypic variation in live mycobacterial cells. As the first proof of concept of our µDeSCRiPTor approach, we used our *M. smegmatis* RecA fluorescent reporter (RecA-GFP)^17^, to screen for compounds that induce RecA fluorescence, aiming to make the population more susceptible to treatment, and decrease fluorescence variation, aiming to homogenize single-cell responses (Fig. 2a). A cytosolic red-fluorescent marker (mCherry_cyt_) was also added for automated segmentation of microcolonies over time (Fig. 2b). To assess the effect of PTC treatment, we computed indices of RecA-GFP mean fluorescence (µ_RecA-GFP_) and fluorescence variance (σ^2^_RecA-GFP_) and estimated the effect sizes for those indices, by comparing the pre-exposure stage with the compound-exposure stage, upon normalization to the negative control group treated with DMSO (Fig. 2b; Methods). Large effect sizes were associated with strong changes in the fluorescence-derived indices upon treatment. An ideal PTC was expected to cause significant increase in the index (µ_RecA-GFP_) and significant decrease in the index (σ^2^_RecA-GFP_) compared to DMSO.

**Figure 2.**
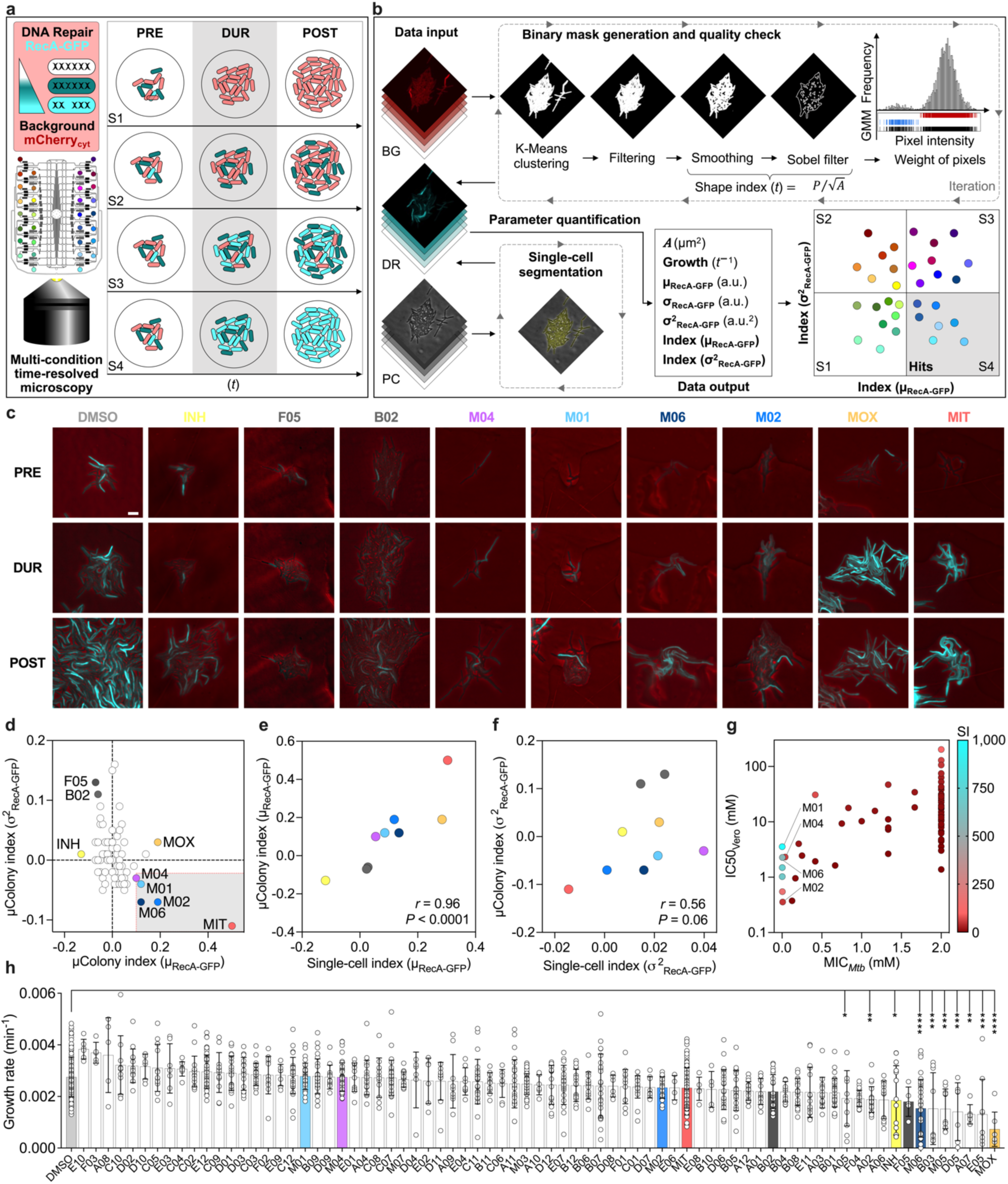
Rationale and outcome of µDeSCRiPTor. **a**,**b**, Schematics of screening setup (**a**) and analysis workflow (**b**). Green-fluorescent reporter of DNA damage response (DR), constitutively expressing a cytosolic red-fluorescent marker as a background (BG). Green gradient denotes the magnitude of response to DNA damage severity. The bacterial suspension is loaded into the 32-condition platform and bacteria are imaged before (PRE), during (DUR), and after (POST) exposure to PTC (gray shading). Four possible responses are depicted (S1 to S4), relative to changes in RecA fluorescence intensity and cell-to-cell variation (**a**). Three channels for BG and DR fluorescence and phase-contrast (PC) are extracted from time-resolved image series. Direction of the analysis workflow is shown by black arrows, while dashed rectangles and gray arrowheads indicate temporal reiteration (**b**). BG fluorescence was used to generate binary masks for whole-colony segmentation (Methods). Microcolony growth rate was calculated based on their area (*A*), DR due to drug exposure was assessed based on fluorescence values, considering both average (µ_RecA-GFP_) and variance (σ^2^_RecA-GFP_). PTC hits fall into S4 (gray shading). Single-cell analysis was carried out manually (Methods). **c**, Time-lapse images of *M. smegmatis* DR-BG reporter at the end of PRE, DUR, and POST stages. DR (cyan) and PC (red) channels are merged. Scale bar, 5 µm. **d**, Size of the PTC effect (effect size) on both average fluorescence (µ_RecA-GFP_) and variance (σ^2^_RecA-GFP_) when comparing DUR versus PRE stage. Estimates were obtained based on 2 to 15 independent experiments (5 < *n* < 86 microcolonies per condition). Negative control DMSO (dashed black lines), positive control MIT (red circle) and 20% of MIT effect (dashed red lines) are shown. PTC hits (gray shading). **e**,**f**, Spearman correlation between single-cell and microcolony indices of RecA-GFP mean intensity (**e**) and variance (**f**), color-coded as in **c** and **d**. **g**, MIC in *M. tuberculosis* and IC50 in Vero cells of different PTC, and selectivity index (SI: IC50/MIC) as a proxy for cytotoxicity. Data are averaged from two independent experiments. **h**, *M. smegmatis* microcolony growth rate during treatment with different compounds. Error bars represent mean ± SD (5 < *n* < 138, from at least two independent experiments). Significance by two-way ANOVA followed by Dunnett’s multiple comparisons test: \**P* < 0.05; \*\**P* < 0.005; \*\*\**P* < 0.0005; \*\*\*\**P* < 0.0001.

We tested two categories of compounds: 65 fragment compounds with millimolar activity^22^; and 7 phenanthroline derivatives with micromolar activity^23^ (Supplementary Table 1). The experimental setup was divided into three stages: 6 hours of pre-growth in standard medium (PRE); 6 hours of exposure to subinhibitory concentrations of candidate PTC or control molecules (DUR); and 6 hours of washing (POST) (Fig. 2a and c; Supplementary Video 2; Source Data 2). The responses of individual microcolonies during the screening were plotted in a two-dimensional graph displaying the relationship between the two indices of fluorescence, corresponding to four possible scenarios (Fig. 2a and d). As a positive control we used MIT, which causes double-strand DNA breaks, resulting in significant induction of index (µ_RecA-GFP_) and significant reduction of index (σ^2^_RecA-GFP_). Thus, we regarded as PTC hits all compounds falling into the lower-right quadrant (Fig. 2b), causing a phenotypic change not less than 20% of the effect caused by MIT (Fig. 2d; Extended Data Figs. 2a and b; Supplementary Table 2). Among them, M02 and M06 caused the strongest and most significant effects in both indices, while M01 and M04 caused significant increase in the index (µ_RecA-GFP_), and moderate but not significant decrease in the index (σ^2^_RecA-GFP_). Most of the tested compounds did not induce major phenotypic changes toward the other three possible scenarios, except for F05 and B02, which caused a significant effect opposite to that of PTC hits. Interestingly, treatment with the anti-tubercular drugs moxifloxacin (MOX) or isoniazid (INH) caused a sharp increase or decrease, respectively, in the index (µ_RecA-GFP_), with high levels of phenotypic variation. We also carried out a separate analysis at the single-cell level, but only for a targeted group of samples due to limited segmentation capacity. We were able to show positive correlations between the results obtained from microcolonies and those obtained from single cells (Figs. 2e and f; Extended Data Figs. 2c−e). However, the correlation between indices (σ^2^_RecA-GFP_) was weaker due to the low number of single cells segmented, especially in the PRE stage (Fig. 2f; Extended Data Fig. 2e).

About 94% of the PTC tested showed a good selectivity index between the half maximal inhibitory concentration (IC50) in Vero cells and the minimum inhibitory concentration (MIC) in the tubercular pathogen, with particularly high values for the selected PTC hits, implying low cellular toxicity and high potency (Fig 2g). However, the principle used to select PTC in *M. smegmatis* was not based on growth inhibition (Fig 2h), but exclusively on RecA pheno-tuning (Fig 2d), and only one PTC hit also affected growth rate. In conclusion, the µDeSCRiPTor strategy enabled us to select four compounds (Extended Data Fig. 2f) that met our expectations for PTC and were further analyzed.

### PTC hits induce DNA repair mechanisms and may weakly interfere with DNA gyrase

To probe the effect of the identified PTC hits on the mycobacterial cell, we carried out whole-transcriptome analysis in *M. tuberculosis*. We investigated the rapid transcriptional remodeling of axenic cultures upon treatment with PTC or MOX (10X MIC) for approximately a quarter of the generation time. Principal Component Analysis (PCA) revealed that biological variability was the main source of variance among our datasets and that three out of four PTC, namely, M01, M02 and M06, clustered together, implying a similar function (Fig. 3a). This was confirmed by differential analysis of gene expression in PTC-relative to DMSO-treated bacteria, as clustered PTC shared 898 differentially expressed genes (DEGs), whereas all four PTC shared 317 DEGs (Figs. 3b; Supplementary Table 3). Consistent with our screening results (Fig. 2), PTC and MOX were found to share 116 DEGs, which were associated with DNA damage response and repair (Fig. 3c; Supplementary Table 4). However, the extent of *recA* transcriptional induction was more pronounced in response to the three clustered PTC as compared to both MOX and M04 (Fig. 3d). M04 also turned out to be quite insoluble, leading us to not prioritize this hit. Thus, we focused on the clustered PTC and especially on M06, which had better physicochemical properties and more promising activity profile in combination with other anti-tubercular drugs (Supplementary Fig. 1a).

**Figure 3.**
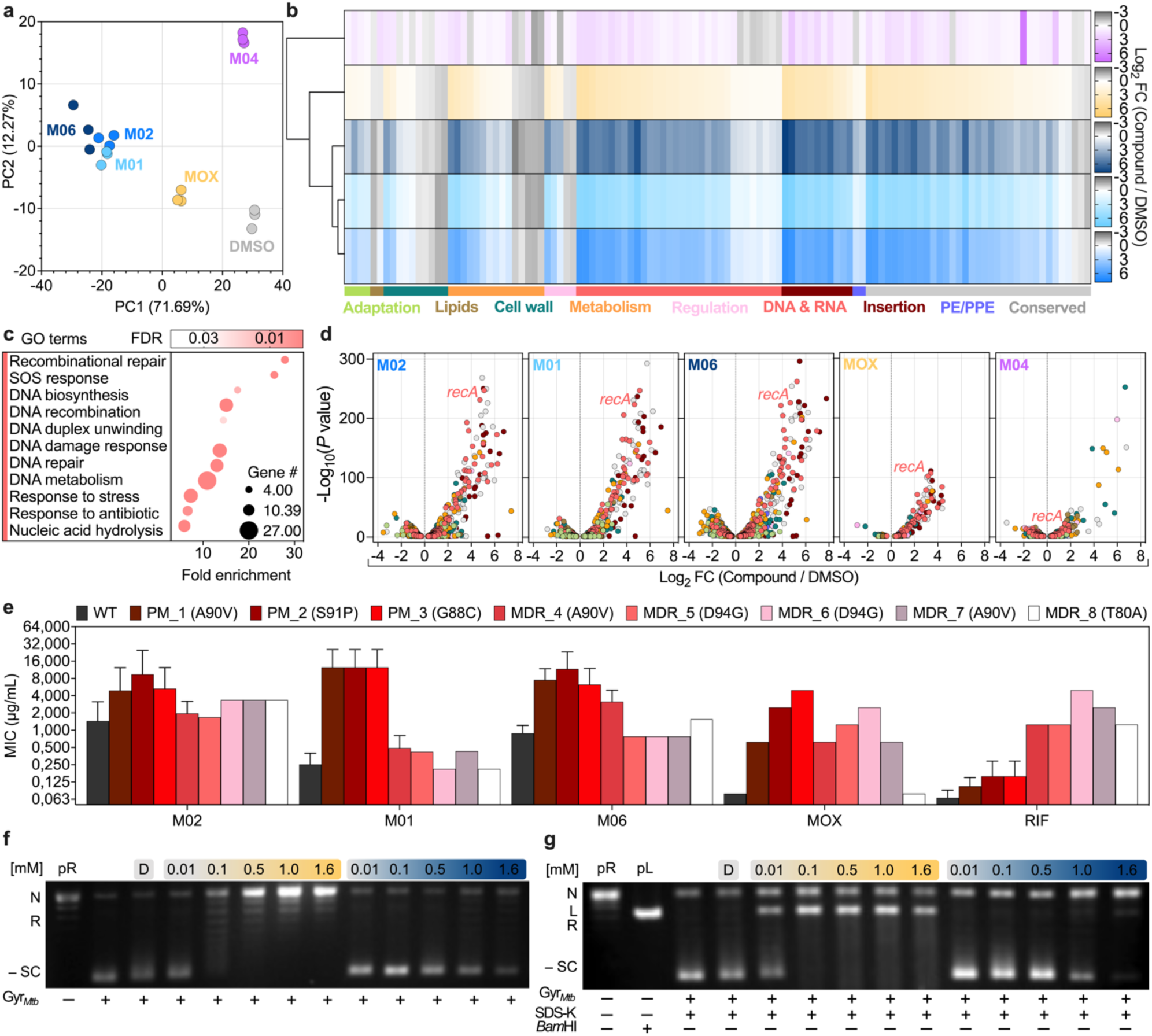
Transcriptome analysis of PTC-treated *M. tuberculosis* and effects on DNA gyrase. **a**, Exploration of experimental variability (*n* = 3) from the first two components of a PCA, with percentages of variance associated with each axis. PC1 (biological variation) describes more than 70% of the total variance in the dataset. **b**, Heat maps showing the log_2_ fold change (FC) of genes that are differentially regulated in compound-treated versus DMSO-treated bacilli (color-coded as in **a**), based on a generalized linear model on the counts. Samples clustering of normalized data is shown. Genes are also grouped into color-coded functional categories. **c**, Gene Ontology (GO) enrichment analysis of 116 shared genes that were differentially regulated upon treatment with PTC hits or MOX compared to DMSO. Complete GO biological processes, including electronic and manually curated annotations, were identified by overrepresentation test against all *M. tuberculosis* genes in PANTHER database. Raw *P*-values were determined by Fisher’s exact test, and False Discovery Rate (FDR < 0.05) was calculated by Benjamini-Hochberg procedure. Bubble chart reports the most specific functional categories sorted by fold enrichment. The full hierarchies of significantly enriched functional categories are also available (Supplementary Table 4). **d**, DEGs in compound-treated compared to DMSO-treated bacilli ranked in volcano plots according to -log_10_ adjusted *P* value as a function of log_2_ FC ratio. Functional categories are color-coded as in **b**. **e**, MIC of clustered PTC and anti-tubercular drugs in *M. tuberculosis* WT, *gyrA* point mutants (PM 1 to 3) and multi-drug resistant strains (MDR 4 to 8). GyrA amino-acid changes are indicated in brackets. Error bars represent mean ± SD (*n* ≥ 2). **f**,**g**, Representative supercoiling (**f**) and cleavage (**g**) assays of relaxed pBR322 (pR) by *M. tuberculosis* DNA gyrase (Gyr*Mtb*) in the absence or presence of DMSO (D) and with increasing concentrations of MOX (orange gradient) and M06 (blue gradient). Nicked (N), relaxed (R), linearized (L), and negatively supercoiled (-SC) DNA is indicated. SDS and proteinase K (SDS-K) were used to denature and remove DNA-bound Gyr*Mtb* and reveal latent DNA breaks (**g**).

PTC hits are heterocyclic organic compounds derived from a phenanthroline scaffold containing two carbonyl groups and resemble the portion of quinolones responsible for binding to DNA gyrase^24^. Thus, we probed whether DNA gyrase could be the target. We first tested a panel of *M. tuberculosis* strains carrying the most frequent *gyrA* mutations conferring moxifloxacin resistance (Supplementary Table 5). While laboratory strains harboring *gyrA* point-mutations were moderately resistant to PTC, especially to M02 and M06, all multi-drug resistant clinical isolates were found to be fully susceptible to PTC (Fig. 3e). Next, we directly tested the effect of M06 against purified *M. tuberculosis* DNA gyrase in vitro. As opposed to MOX, which caused strong inhibition of the negative supercoiling activity of DNA gyrase as expected, M06 caused very weak inhibition of DNA gyrase at a ten-fold higher concentration (Fig. 3f). Likewise, MOX promoted plasmid DNA breaks via stabilization of gyrase-DNA cleavage complex starting from low µM concentration, whereas M06 causes limited DNA rupture only at mM concentration (Fig. 3g). Additionally, docking analysis predicted that M06 could occupy the active site of DNA gyrase, interacting via hydrogen bond of the hydroxyl group (O4) and hydrophobic contact (C14) with the residue D94, which is in contact with the magnesium ion^24^ (Supplementary Figs. 1b−e).

PTC hits also caused a broader transcriptional response different from that of MOX (Extended Data Figs. 3a−e; Supplementary Table 4). Specifically, the three clustered PTC caused marked repression of the electron transport chain, cellular respiration, glycolysis, and tricarboxylic acid cycle, as well as of fatty acid and lipid metabolism (Extended Data Figs. 3c−e). Overall, these results are suggestive of a broader and severe mode of action of PTC against the mycobacterial cell.

### PTC mechanism of action is associated with the arylamine N-acetyltransferase

To further clarify the mechanism of action, we searched for mutants resistant to the PTC hits, and isolated two on M06 and two on M02 (Supplementary Table 6). Interestingly, three out of four shared the same mutation on the *rv3566c* gene (Fig. 4a), which encodes the arylamine N-acetyltransferase (NAT). NAT is a highly conserved xenobiotic metabolizing enzyme, which uses acyl donors, such as acetyl-CoA or propionyl-CoA, to acetylate diverse heterocyclic amines^25^. For instance, NAT was shown to antagonize the INH-activating role of the catalase peroxidase KatG (Extended Data Fig. 4a), by acetylating this prodrug^26^. Additionally, *M. tuberculosis rv3566c* is located downstream of an operon implicated in lipid catabolism (Extended Data Fig. 4a) and was associated with mycolic acids biosynthesis, cell-wall integrity and intracellular survival^25, 27^. Therefore, we sought to clarify whether NAT was responsible for the inactivation or the activation of PTC or if it was the target. In all three isolated mutants, NAT was devoid of leucine 81 (Fig. 4a), which is implicated in enzyme structure stability^28^, and is located within the active site between two critical catalytic residues (Extended Data Fig. 4b). NAT mutants did not experience any growth defect (Fig. 4b) but were cross-resistant to the clustered PTC (Fig. 4c). This was consistent with transcriptional decrease of *nat* in PTC-stressed bacilli (Supplementary Table 4). To further probe the role of NAT into the mechanism of action of PTC, we generated an inducible knock-down (*si_nat*) and an overexpressing (*oe_nat*) *M. tuberculosis* strain (Extended Data Figs. 4c and d). Transcriptional repression of *nat* caused resistance to the three clustered PTC (Fig. 4d), supporting the hypothesis that NAT is the cellular target, and not the inactivating enzyme. In contrast, transcriptional induction of *nat* did not cause PTC resistance (Fig. 4e), implying a more complex mechanism^29^.

**Figure 4.**
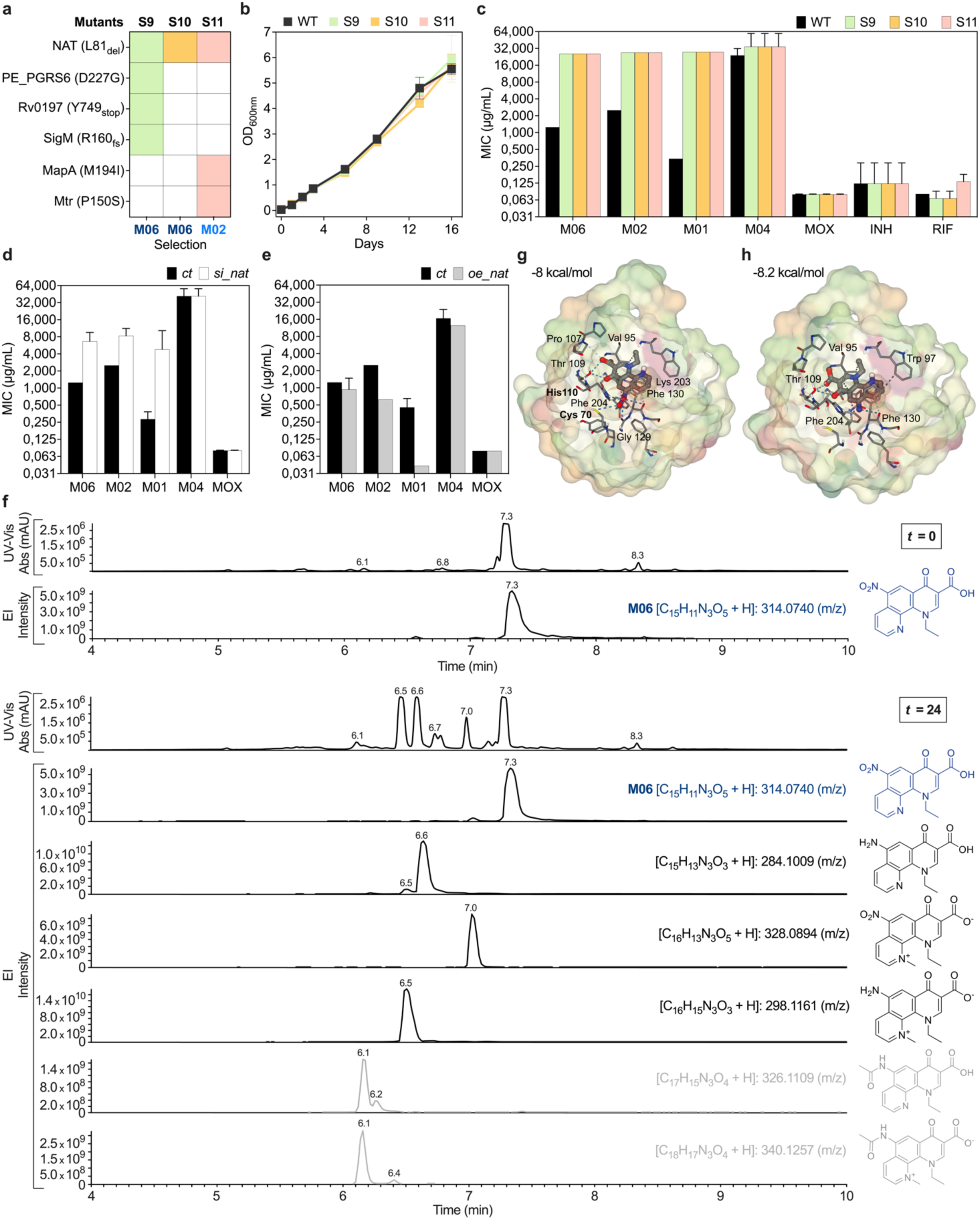
Characterization of the mechanism of action of clustered PTC. **a**, Summary of variant-calling analysis of three spontaneous mutants sharing the same mutation in *rv3566c* gene (*nat*). The compound on which the mutants were isolated is indicated. **b**, Growth kinetics of *M. tuberculosis* wild type (WT), S9, S10, and S11 mutants. Error bars represent mean ± SD (*n* = 2). **c–e**, MIC of PTC and anti-tubercular drugs in *M. tuberculosis* WT and *nat* mutants (**c**); following silencing of *nat* (*si_nat*) (**d**); and overexpression of *nat* (*oe_nat*) (**e**) and in related controls (*ct*). Error bars represent mean ± SD (*n* = 3). **f**, Representative spectra at 270 nm (UV) and extracted ion (EI) chromatograms of supernatant from *M. tuberculosis* treated with M06 (15X MIC) at the onset of treatment and after 24 hours. X-axes indicate time after injection (min). Y-axes indicate intensities (a.u.). Chemical structures, formulas, and mass-to-charge ratio of M06 and its metabolites are shown next to the relevant peaks (*n* = 2). **g**,**h**, In silico docking poses of CPK-colored M06 (**g**) and its amino derivative (**h**) in the catalytic pocket of NAT. Binding affinity is indicated (kcal/mol). Predicted amino acid residues in hydrophobic contact (dark gray) and linked by either weak (light gray) or strong (blue) hydrogen bonds with ligands atoms are shown. Bolded residues are part of the catalytic triad (**g**).

To determine whether NAT is the target or the enzyme that activates the top PTC hit M06, contingent on reduction of the nitro group at position N3 (Supplementary Fig. 1b), we carried out liquid chromatography and mass spectrometry analysis of the supernatant of *M. tuberculosis* treated with M06. After 24 hours of M06 treatment, we found more than a quarter of the original molecule and three major metabolites: about 10% methylated at position N1; 28% nitro-reduced at position N3; and 30% of methylated and nitro-reduced variants (Fig. 4f). Importantly, we found only a negligible fraction of acetylated form at position N3, supporting the assumption that M06 is the inhibitor of NAT and not its substrate. Finally, we carried out docking analysis of M06 and its nitro-reduced variant with NAT^30^. Similar to the INH substrate (Extended Data Fig. 4e), both M06 and its reduced metabolite fell into the active site of NAT (Figs. 4g and h). However, while M06 was predicted to bind two of the three catalytic residues (Fig. 4g), its amino derivative was found to lose these interactions (Fig. 4h), supporting that this metabolite is not a substrate of NAT. Collectively, these results led us to conclude that NAT is targeted by M06.

### M06 affects the composition of the mycobacterial cell envelope and causes oxidative stress

Given the substantial transcriptional remodeling related to energy and lipid metabolism triggered by M06 (Extended Data Fig. 3e) and the implication of NAT in lipid metabolism^27^, we probed whether M06 could alter the composition of the cell envelope in *M. tuberculosis*. We fed bacilli with radiolabeled acetate and analyzed fatty acids and lipids by thin layer chromatography (TLC). We found no differences in the TLC profiles of strains expressing wild type or mutated NAT variants, or overexpressing NAT, suggesting that deletion of leucine 81 does not affect the physiological role of NAT (Supplementary Figs. 2a−d). High concentrations of M06 caused significant decrease in mycolic acids, especially of the keto form in all three strains (Figs. 5a and b). These changes did not occur under MOX treatment, while, as expected, INH caused marked inhibition of all forms of mycolic acids and accumulation of fatty acids (Extended Data Fig. 5a). TLC analysis of extractable lipids revealed moderate accumulation of trehalose monomycolate (TMM), which was less pronounced in NAT-mutant strain (Fig. 5c). Slight accumulation of TMM was also observed after treatment with MOX (Extended Data Figs. 5b and c). Lastly, at high concentrations of M06 we found significant decrease in phosphatidylethanolamine (PE), phosphatidylinositol (PI), phoshpatidylinositol mannosides (PIM), as well as trehalose dimycolate (TDM) and TMM (Fig. 5c; Extended Data Figs. 5b and c). These results were also confirmed at the single-cell level, as bacilli treated with M06 exhibited decreased staining of cell-wall lipids, except if NAT was mutated (Figs. 5d and e).

**Figure 5.**
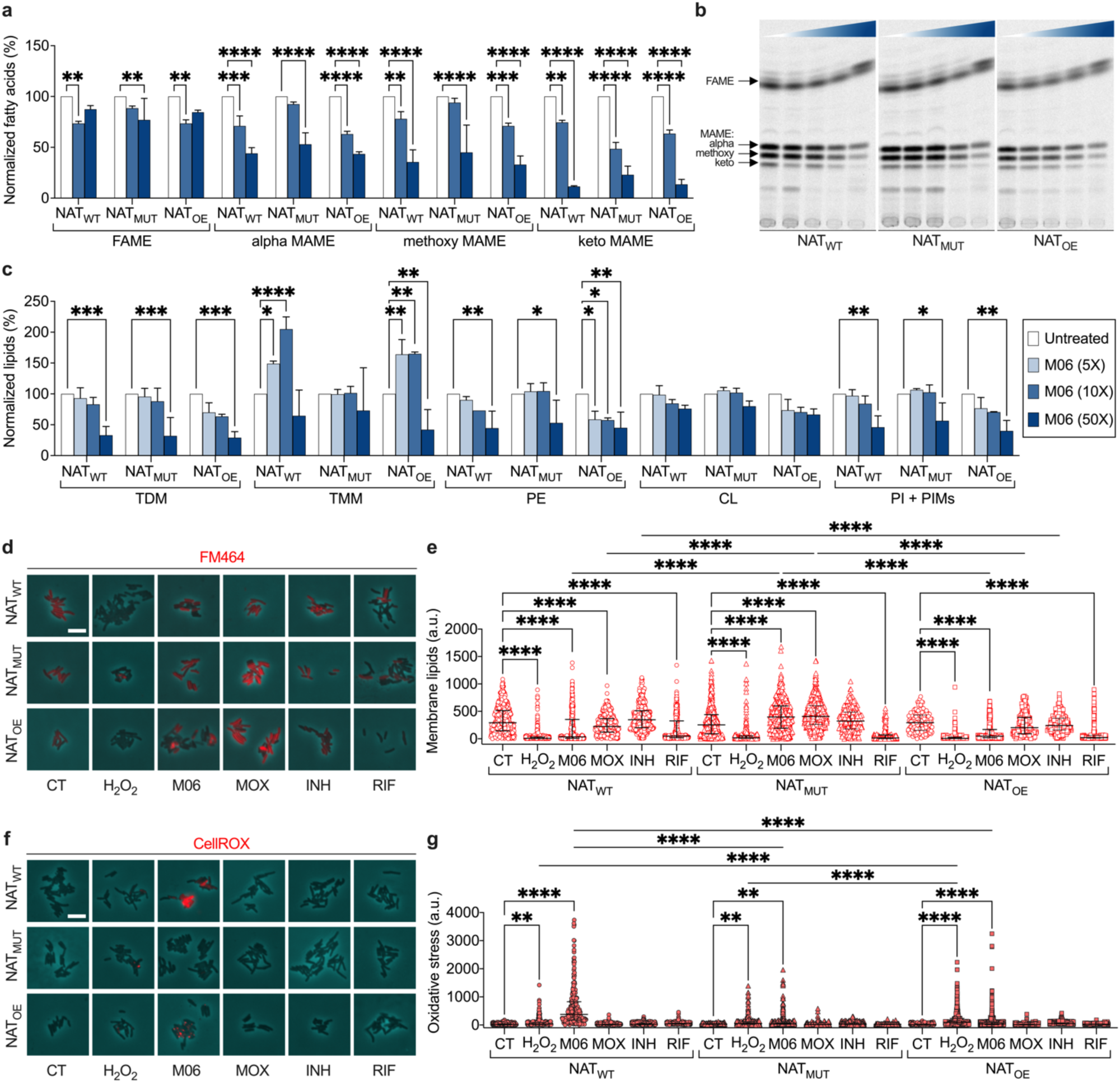
Lipid profiling and analysis of oxidative stress upon M06 treatment. **a**,**c**, Quantification of fatty acid methyl esters (FAME) and different types of mycolic acid methyl esters (MAME) (**a**), and of trehalose esters of mycolates (TDM; TMM), phospholipids (PE; CL), and population of PI and PIMs (**c**) in exponentially growing *M. tuberculosis* WT (NAT_WT_); S10 mutant (NAT_MUT_); and NAT overexpressing strain (NAT_OE_), treated with different concentrations of M06 relative to the MIC for 24 hours (inset). Data are normalized to untreated samples and error bars represent mean ± SD (*n* = 2). Significance by two-way ANOVA followed by Dunnett’s multiple comparisons test: \**P* < 0.05; \*\**P* < 0.01; \*\*\**P* < 0.001; \*\*\*\**P* < 0.0001. **b**, Representative TLC of extracted FAME and MAME (arrows). Color gradient indicates from left to right the absence or increasing concentrations of M06 equal to 5X, 10X, 25X and 50X MIC (1.25 µg/mL). **d**,**f**, Representative snapshot images of exponentially growing *M. tuberculosis* strains (as in **a**,**c**) exposed for 24 hours to DMSO (CT), H_2_O_2_ (30 mM) or to different drugs (10X MIC). Bacteria were stained for membrane lipids (**d**) or oxidative-stress (**f**) and imaged by phase contrast (cyan) and fluorescence (red). Fluorescence images are scaled to the brightest frame. Scale bars, 5 μm. **e**, Single-cell snapshot analysis of FM464 fluorescence from strains and conditions in (**d**). Data are from 2 independent experiments (160 < *n* < 600 single cells per condition). Black lines indicate median with interquartile range. Significance by two-way ANOVA followed by Tukey’s multiple comparisons test: \*\*\*\**P* < 0.0001. **g**, Single-cell snapshot analysis of CellROX fluorescence from strains and conditions in (**f**). Data are from 2 independent experiments (200 < *n* < 850 single cells per condition). Black lines indicate median with interquartile range. Significance by two-way ANOVA followed by Tukey’s multiple comparisons test: \*\**P* < 0.005; \*\*\*\**P* < 0.0001.

Cell-wall alterations can result from both inhibition of synthesis and direct degradation. Interestingly, transcriptomic analysis revealed significant deregulation of genes involved in detoxification reactions upon M06 treatment (Supplementary Table 4). We further confirmed this by targeted gene expression analysis and showed that M06 causes deregulation of genes responsible for antioxidant defense, including increased *katG*, which protects against reactive oxygen and nitrogen intermediates; decreased *trxB1*, important for redox balance; and decreased *ahpE*, *ephA* and *ephD*, involved in detoxification from organic peroxides and rescue of oxidized lipids (Extended data Fig. 5d). Conversely, NAT mutant strains treated with M06 maintained unchanged levels of genes related to detoxification from lipid peroxidation, implying their lower susceptibility to this PTC. To check whether M06 induced oxidative stress, we carried out single-cell snapshot analysis. We confirmed that *M. tuberculosis* exposed to M06 shows significant levels of oxidative stress, which was not the case for moxifloxacin or other anti-tubercular drugs tested and was less pronounced in both NAT mutant and NAT overexpressing strain (Figs. 5f and g). Overall, these results imply that M06 affects the integrity of mycobacterial cell envelope by both directly targeting NAT and inducing oxidative stress.

### The top PTC M06 shows promise in potentiating existing anti-tubercular drugs

To examine whether pheno-tuning RecA expression made *M. tuberculosis* more sensitive to treatment, we tested the activity of our top PTC M06 in combination with some standard antitubercular drugs^11^. We combined the highest concentration of M06 that had no bactericidal action per se (Supplementary Fig. 3a) with a low concentration of a second drug. Pharmacodynamics were estimated via colony forming units (CFU) quantification for a minimum of two weeks up to a maximum of six weeks, or until growth rebound was observed. Among the first-line drugs, we found that M06 had no enhancing effect on ethambutol (EMB), despite what could have been inferred from FIC-index estimation (Extended Data Fig. 6a; Supplementary Fig. 1a). As expected for a drug that inhibits NAT, the association of INH with M06 decreased bacterial survival by 10 to 100 folds compared to INH alone (Fig. 6a). The strongest enhancing effect of M06 occurred in combination with rifampicin (RIF), causing about 30-fold reduction in bacterial survival after a week of treatment compared with RIF alone, down to below the detection limit at two weeks of treatment (Fig. 6b). Additionally, the lack of growth rebound implied complete sterilization of the axenic culture. Consistent with the weak inhibition of DNA gyrase by M06 (Fig. 3), we also found moderate additive effect of M06 in combination with the second-line drug MOX (Extended Data Fig. 6b). Interestingly, M06 was also found to decrease the expression of *mfpA* (Supplementary Fig. 3b; Supplementary Table 4), which protects DNA gyrase from fluoroquinolones^31^.

**Figure 6.**
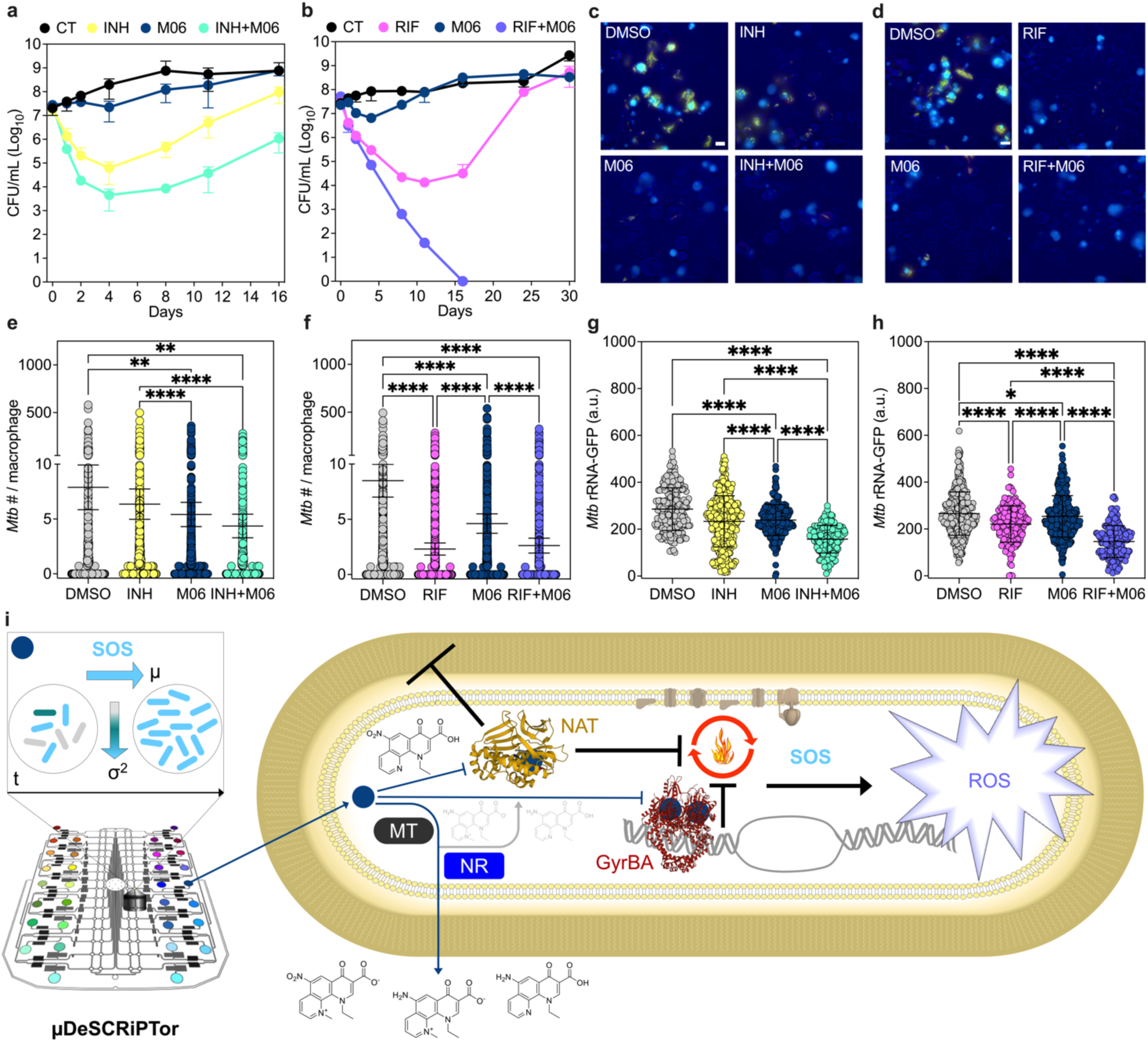
Extracellular and intracellular activity of M06 in combination with first-line antitubercular drugs. **a**,**b**, Drug activity in exponential-phase *M. tuberculosis* cultures grown in the absence of drugs (CT), or in the presence of M06 alone (1X MIC), of individual anti-tubercular drugs alone (2X MIC), or of their combinations: INH ± M06 (**a**); RIF± M06 (**b**). CFU are expressed as mean ± SD (2 ≤ *n* ≤ 3). **c,d**, Representative snapshot images of RAW 264.7 macrophages at 6 days post infection with *M. tuberculosis* rRNA-GFP_DsRed2^32^ (*n* = 6). Macrophages were left untreated (DMSO) and treated for 6 days with INH ± M06 (**c**) at the concentrations used in (**a**), or with RIF ± M06 (**d**) at the concentrations used in (**b**). Bright field (blue), rRNA-GFP (green), DsRed2 (red), and DRQ7 (cyan) fluorescence are merged. Scale bars, 10 μm. **e**,**f**, Number of intracellular bacilli at 6 days post infection, left untreated (DMSO) and treated with INH ± M06 (**e**), or with RIF ± M06 (**f**). The number of bacilli was estimated from the area of the clump divided by the average size of a single bacillus. Error bars represent mean ± confidence interval (*n* = 6): 1236 ≤ macrophages ≤ 2097 per condition (**e**); 2022 ≤ macrophages ≤ 3177 per condition (**f**). Significance by Kruskal-Wallis one-way ANOVA: \*\**P* < 0.005; \*\*\*\**P* < 0.0001. **g**,**h**, rRNA-GFP fluorescence of intracellular bacilli at 6 days post infection in untreated macrophages (DMSO) or macrophages treated with INH ± M06 (**g**), or with RIF ± M06 (**h**). Error bars represent mean ± SD from 6 independent experiments (**g**, 345 ≤ bacilli ≤ 521) and (**h**, 276 ≤ bacilli ≤ 627). Significance by two-way ANOVA followed by Tukey’s multiple comparisons test: \**P* < 0.05; \*\*\*\**P* < 0.0001. **i**, Schematic of the dynamic microfluidic single-cell screening for PTC, based on SOS induction and decreased cell-to-cell variation, and of the expected mode of action of the main PTC hit (blue circle). Blue sharp arrows indicate entry of M06 and exit of its main metabolites (structures are shown). Blue blunt arrows indicate targets inhibition by M06. Gray arrow and structures indicate unlikely substrates of NAT upon M06 bioprocessing by methyltransferases (MT) and nitroreductases (NR). Black blunt arrows indicate resulting inhibition of cell-envelope and metabolic processes. Black sharp arrow indicates resulting induction of reactive oxygen species (ROS), which in turn affect cellular integrity.

Next, we tested the effect of M06 in combination with INH and RIF intracellularly. We infected macrophages with a strain of *M. tuberculosis* expressing GFP from the native ribosomal promoter, to assess the metabolic activity, and constitutively expressing DsRed2^32^ (Figs. 6c and d). At six days post infection, we found that M06 alone causes significant decrease of the bacterial burden (Figs. 6e and f), consistent with the essentiality of NAT in macrophages^27^. Additionally, the combination of M06 with INH significantly reduced the fraction of infected macrophages and the relative bacterial load compared to INH alone (Fig. 6e; Extended Data Fig. 6c). Intracellular bacilli also showed decreased metabolic activity in the presence of M06 alone, which was more pronounced in combination with INH, as compared to INH alone (Fig. 6g). However, the M06-INH combination increased macrophage mortality (Extended Data Fig. 6d). The combination of M06 with RIF significantly reduced the fraction of infected macrophages compared to RIF alone, with a similar impact on the bacterial burden per macrophage (Fig. 6f; Extended Data Fig. 6e). This might hinge on the fact that M06-RIF combination becomes increasingly effective in vitro only after six days of exposure (Fig. 6b). We also found that M06-RIF combination significantly decreases *M. tuberculosis* metabolic activity (Fig. 6h), as well as macrophage mortality (Extended Data Fig. 6f).

Collectively, these results prove that the µDeSCRiPTor strategy successfully identified a pheno-tuning and drug-potentiating compound, undermining the tubercular pathogen via a complex mode of action (Fig. 6i).

## Discussion

The discovery of new effective antimicrobials is a complex and inefficient process in need of renewal^7, 9, 15^. Despite the critical role of microbial phenotypic variation in treatment failure, this phenomenon is hardly included in conventional models of drug discovery due to technical hindrances, significantly limiting their potential^2, 6, 12, 14^. To address this issue, we conceived the µDeSCRiPTor strategy, which is based on multi-condition microfluidics, to dynamically capture phenotypic changes via spatiotemporal imaging of individual cells within microcolonies. We proved the full potential of our strategy on mycobacterial cells, based on our earlier finding that pre-existing phenotypic variation in DNA damage response is associated with differential drug susceptibility at the subpopulation level^17^. However, this approach can be virtually applied to any instance of phenotypic variation that is critical to a given microbial pathogen to endure^5, 6^, aiming to induce a homogeneous state of hypersensitivity. Although our initial goal was a proof of concept to probe changes in a fluorescent reporter upon exposure to different compounds, we were already able to identify a promising PTC that deserves further consideration for optimization and drug development. The advantage offered by microfluidic technologies for studying the dynamics of single-cell behaviors comes either at the expense of throughput or at the expense of resolution and flexibility^18^. To fill these gaps, we customized a multi-condition microfluidic platform. Its peculiarity lies in the combination of continuous flow, typical of passive devices, with pneumatic valves, typical of active devices^19^. Given the atypical aspect ratio of the microchambers featured in this platform, they can be actuated with pressures significantly lower than those used for on-off micromechanical valves^33^, generating a gentle pneumatic cell-trapping mechanism, compatible with flow of nutrients and cell viability. Overall, this platform allowed us to maintain high spatiotemporal resolution of single-cell behaviors and maximum control over environmental dynamics, as well as to increase the experimental throughput. However, the microfabrication process of this PDMS-based prototype includes several manual steps and currently has a success rate of about 60-70%, mainly due to adhesion defects between different layers that can cause fluid leaks. Additionally, the percentage of bacterial cells growing averaged 76%, about 10% less than for simpler devices^17^. This was most likely due patchy adhesion of the flexible PDMS membrane to glass, retarding access to nutrients. Thus, we had to reject all videos containing bacteria that were not growing from the beginning of the experiment, that were growing in 3D, or moving due to membrane instability, and those with too many out-of-focus frames. Taking that into account, the overall success rate of the whole screening procedure averaged 54%. This rate can be improved by modifying the existing prototype with new materials and the fabrication procedure, such as implementing hot embossing or additive manufacturing of hard thermoplastics and elastomers^34^. Upgrading this platform may also involve increasing the number of conditions that can be simultaneously tested in a more compact and robust device. Despite these manufacturing hindrances, we accomplished a first screening, and identified four initial PTC hits that induced more homogeneous activation of RecA, as a proxy of the SOS response^21^.

To assess the effects of our candidate PTC on the SOS response within clonal mycobacterial cells, we relied on standardized effect-size calculation and their accuracy was estimated with confidence intervals^35^. Mixed-effects models were used to estimate effect sizes on two fluorescence indices. Such model-based approaches are suited to accommodate various technical effects as well as missing values due to out-of-focus images^36^. In order to control technical variation across experiments we advise to add either a passive reference dye such as FITC used in our study, or fluorescent microbeads in all experimental replicates. We considered as biologically relevant all effects equal to at least one-fifth of that caused by the control agent MIT. However, this value is to be determined on a case-by-case basis. The present case resulted in four initial PTC hits, which would have been overlooked by standard selection criteria^7, 9, 15^. Unlike selections dictated solely by macroscopic population-averaged effects, the key advantage of the µDeSCRiPTor approach is that it allows us to visualize small dynamic changes on individual cells that would be unattainable by static or even dynamic bulk-cell methods. Furthermore, it allows us to identify subtle differences between compounds clustering in the same functional class that otherwise would not be appreciated. Nevertheless, the reported strategy has a throughput of at least one or two orders of magnitude lower than standard drug discovery approaches^9, 15^, mainly due to manual manufacturing of the platform, long experimental duration and limited automated analysis, which could be improved by implementing deep neural network segmentation tools^37^. Therefore, one possibility is to preselect compounds based on other parameters, without necessarily increasing the likelihood of identifying a PTC. For instance, our candidate PTC included primarily small molecule compounds with no apparent activity^22^, and a few quinolone analogs^23^, which were the source of our hits. However, for a deeper characterization, we focused on the PTC that had not only the strongest effect on the SOS indices but was also the most soluble.

Indeed, M06 proved particularly attractive from both a mechanistic and an activity point of view, confirming the soundness of the µDeSCRiPTor approach in homogenizing clonal cells to make them more susceptible to drug-mediated killing. Similarly, it was proposed that noise modulators in HIV^38^ and genetically collapsing phenotypic variation in a tuberculosis model^39^ may improve treatment outcome. Here we propose that the main target of our top PTC is NAT, whose expression levels fall upon M06 treatment, and whose genetic modification of the active site and transcriptional inhibition cause M06 resistance. NAT is a ubiquitous enzyme involved in N-acetylation of xenobiotics, causing either substrate inactivation or activation^25, 26, 28^. However, the main metabolization processes that M06 undergoes in the mycobacterial cell are nitroreduction and methylation, occurring either alone or in combination, likely inactivating the original compound^23^. This is not surprising since *M. tuberculosis* genome encodes several nitroreductases and methyltransferases, responsible for either prodrug activation or drug inactivation^40, 41^. In the case of M06, NAT-mediated N-acetylation could take place only following reduction of the nitro group with the formation of an arylamine, which is one of the possible substrates of NAT^28^. However, we found that the levels of acetylated M06 are negligible, ruling out that M06 is a NAT-activated prodrug. Moreover, other N-acetyltransferases in the mycobacterial cell could be responsible for this modification^42^.

NAT was already reported to be a promising anti-tubercular target, as strains devoid of this enzyme show defects in cell-wall mycolates and associated complex lipids, and reduced survival in macrophages^27^. Given the divergence between the human and mycobacterial NAT enzymes, piperidinol analogs, heterocyclic compounds different from our PTC, were formerly identified as potent selective inhibitors of mycobacterial NAT^43^. In support of the assumption that M06 targets NAT, we show that M06 treatment causes decrease in several cell-envelope components, particularly ketomycolates, which are implicated in cell-wall stability, biofilm formation and drug tolerance^44^. The alterations we observed recall, albeit on a much smaller scale, the inhibition of fatty acid synthase II by INH, which directly targets mycolic acids biosynthesis. Importantly, the reduction in ketomycolates upon M06 exposure was not associated with an accumulation of their precursor hydroxymycolate, which typically hinges on depletion of the epoxide hydrolase EphD^45^. However, we did find that the expression levels of *ephD* decrease in M06-treated *M. tuberculosis* but not in NAT mutant strains. Overall, we speculate that the broad but moderate inhibitory effect of M06 against mycolic acids and other cell-envelope constituents is not due to direct inhibition of biosynthetic pathways, but most likely to indirect causes, including perturbation of acetyl-CoA homeostasis, following NAT inhibition^27, 30^; metabolic remodeling; and oxidative stress. Interestingly, inhibition of cell-wall biosynthesis and cell division was found to trigger the SOS response in *Escherichia coli*^46^. Consistent with this, M06 exposure induced antioxidant defenses, and hindered cell wall biogenesis, energy metabolism and respiration, presumably to attenuate the burden derived from oxidative damage^47, 48^.

The presence of oxidative stress in *M. tuberculosis* treated with M06 may originate from moderate inhibition of DNA gyrase. Indeed, M06 appears to act as an inefficient quinolone^49^, although it both inhibits the catalytic activity of the enzyme and injures DNA. Interestingly, high concentrations of quinolones were shown to cause a paradoxical effect, whereby higher bacterial survival is associated with lower levels of reactive oxygen species and vice versa^50^. We speculate that the weak inhibition of DNA gyrase by M06 may be advantageous for this molecule and lead to lethal levels of ROS, which intoxicate the cell^47, 48^. On the other hand, bioreduction of nitroaromatic compounds has been implicated in bacterial cell poisoning^51^. Thus, the nitro group of M06 may also act as a functional electron trap, contributing to the redox toxicity of this compound in the mycobacterial cell, which can carry out several bioreactions on this PTC^40–42^. Deregulation of redox balance was also shown to alter the activity of human NAT, via reversible oxidation of the catalytic cysteine^52^. Since *M. tuberculosis* NAT mutants are less susceptible to M06 and exhibit significantly lower levels of oxidative stress, at present we cannot exclude that oxidative stress is the primary cause of the anti-mycobacterial activity of M06, and that the mutation found in NAT active site may help the cell to counter this redox imbalance. Only structural analyses will allow us to confirm or disprove the present model, whereby M06 acts via a complex mechanism, which includes two cellular targets, metabolic and redox perturbations, and cellular intoxication with harmful consequences on critical macromolecules (Fig. 6i).

Although M06 per se proved to be an interesting compound, potent in vitro and active intracellularly, the goal of our µDeSCRiPTor strategy was to make cells more homogeneously susceptible to existing therapy. After testing a sample of anti-tubercular drugs, we showed additivity of M06 with INH. This was expected since NAT, enzyme that inactivates INH^26, 28^, is one of the two putative targets of M06. Furthermore, we showed additivity with MOX, presumably because treatment with M06 induces downregulation of MfpA, a protein that protects DNA gyrase from quinolones by mimicking DNA^31^. Finally, M06 strongly synergizes with the RNA-polymerase inhibitor RIF, leading to complete elimination of the persistent fraction in vitro. This might have different explanations. Steric inhibition of the replication fork by M06 in complex with DNA and DNA gyrase could affect not only DNA replication but also the recruitment of RNA polymerase and initiation of transcription^31, 50^. Excessive accumulation of reactive oxygen species upon M06 exposure could make bacilli persistent to RIF increasingly vulnerable^6^. Concerted impairment of cell-wall lipids, which we observe following both M06 and RIF treatment, could fatally weaken the stability of the mycobacterial cell structure. Lastly, acetylation is a conserved post-translational modification critical in all life domains, although its endogenous roles in bacteria remain largely unknown^53, 54^. Interestingly, lysine acetylation of a housekeeping sigma factor was found to promote the activity of the RNA polymerase in actinobacteria^55^. Thus, we cannot exclude that a defect in NAT might have unexpected consequences on transcription as well as on other essential cellular functions.

In conclusion, we developed the µDeSCRiPTor strategy and successfully implemented it with one of the most challenging microbial pathogens, identifying a PTC that potentiates existing anti-tubercular drugs. We also confirmed that phenotypic variation is a promising bacterial target, which can suggest original eradication strategies. We expect that this approach will prove useful to identify other compounds that can enhance antimicrobial therapy, helping us to fight infections that are becoming virtually incurable.

## Methods

### Bacterial strains

Cloning and sequencing were carried out in chemically competent *Escherichia coli* TOP10, grown in LB medium in the presence of appropriate selection: 50 µg/mL kanamycin or 100 µg/mL hygromycin. Mycobacteria were cultured in Middlebrook 7H9 broth supplemented with 0.5% BSA, 0.2% glucose, 0.085% NaCl, 0.5% glycerol and 0.01% Tyloxapol, which was removed for MIC assessment, checkerboard titration and time-lapse microscopy. Middlebrook 7H10 agar was enriched with 10% OADC and 0.5% glycerol. Mycobacterial transformants and primary cultures were selected on 20 µg/mL kanamycin or 50 µg/mL hygromycin, as appropriate. Bacterial stocks were prepared from single colonies grown up to OD_600nm_ 1.0 in the presence of appropriate selection if needed, supplemented with 15% glycerol, and directly frozen at −80 °C. Each aliquot was used only once to start primary cultures. Primary cultures of *M. smegmatis* mc^2^155 (ATCC 700084), *M. tuberculosis Erdman* (ATCC 35801), and *M. tuberculosis H37Rv* (ATCC 27294) were cultured at 37 °C under shacking conditions, at 150 RPM for the fast-growing strain and 50 RPM for slow-growing pathogenic strains, until reaching mid-log phase. For final assays, secondary cultures were obtained by diluting primary cultures 100 times in appropriate medium until mid-log phase was reached.

### Cell lines

RAW 264.7 macrophages (ATCC TIB-71) and Vero cells (ATCC CCL-81) were cultured in Dulbecco’s Modified Eagle Medium (DMEM, Gibco) supplemented with 10% fetal bovine serum (Gibco) and 1X penicillin-streptomycin mix (Gibco). A stock of 10^6^ cells per mL in DMEM and 10% DMSO was defrosted at 37 °C, washed once in 10 mL and inoculated in 30 mL of prewarmed complete DMEM using a T-175 flask. Cells were propagated at 37 °C in humidified 5% CO_2_ atmosphere until confluence. For final experiments, confluent cells were diluted to a concentration between 10^5^ and 2.5*10^5^ cells per mL in complete DMEM without antibiotics.

### Strains construction

Strains, plasmids, and oligonucleotides are listed in Supplementary Table 7. The NAT-overexpressing plasmid pGM321 was constructed by PCR amplifying the *rv3566c* open reading frame, preceded by a Shine Dalgarno sequence optimized for mycobacteria and flanked by *Pac*I and *Hpa*I restriction sites, and cloning it inside the L5 integrative plasmid pND200, under the control of a strong promoter. The ATC-inducible dCas9 *nat* knock-down plasmid pGM315 was constructed by inserting a small guide RNA complementary to the *rv3566c* gene in the L5 integrative plasmid pGM309. Final constructs were electroporated into *M. tuberculosis H37Rv*, and transformants were obtained under hygromycin selection.

### Growth assay

*M. tuberculosis* primary cultures were grown up to OD_600nm_ 0.5-0.6 and diluted to OD_600nm_ 0.025 using pre-warmed Middlebrook 7H9 broth. Growth kinetics were assessed by measuring the OD_600nm_ over a two-week period, in triplicate.

### Chemistry

The PTC candidates used in this study are listed in Supplementary Table 1. Fragment compounds were obtained following the deconvolution of the 6 best hits from a curated small-molecule fragment library of 1604 fragments, combined into 169 pools, which were formerly tested on *M. smegmatis* and *M. tuberculosis*^22^. The 65 singletons derived from this primary screening were purchased from Key Organics UK and showed low inhibition against both mycobacterial species, based on MIC evaluation. The 7 phenanthroline derivatives were selected from a library of 28 compounds synthesized at CERMN^23^ and based on their antimycobacterial activity. See also Supplementary Methods.

### Microfabrication of the 32-condition platform

The designs of the flow and control layer were generated on AutoCAD 2017 (Source Data 1), and corresponding low-resolution plastic photomasks were printed at Selba S.A. The micropatterns were transferred from the photomasks to 100-mm silicon wafers by photolithography, producing one wafer for the flow layer (FL-W) and a second wafer for the control layer (CK-W).

Two silicon wafers were dehydrated at 200 °C on a hot plate for 30 min. The FL-W was produced by spin-coating SU8-2025 photoresist at 3000 RPM for 30 sec, to generate a layer of 30 µm, and soft-baking at 65 °C for 1 min, followed by an incubation at 95 °C for 5.5 min. The photoresist was crosslinked under 155 mJ/cm^2^ UV light for 11.5 sec, and the wafer was baked at 65 °C for 1 min, followed by 95 °C for 5 min. Finally, the FL-W was developed by immersion into propylene glycol methyl ether acetate (PGMEA) for 4 minutes and washed extensively with isopropanol. The CL-W was generated by spin-coating the SU8-2100 photoresist at 1500 RPM for 30 sec, to generate a layer of 200 µm, and soft-baking at 65 °C for 5 min, followed by 95 °C for 20 min. The photoresist was crosslinked under 240 mJ/cm^2^ UV light for 21.8 sec, and the wafer baked at 65 °C for 5 min, followed by 95 °C for 10 min. Finally, the CL-W was developed by immersion into PGMEA for 15 minutes and washed extensively with isopropanol. The FL-W and CL-W were further hard-baked at 180 °C for 2 hours, and silanized with Trichloro (1H,1H,2H,2H-perfluorooctyl) silane in a vacuum chamber overnight. The microchannels height was profiled with a DektakXT (Bruker).

For soft-lythography, the FL-W was covered with a degassed mixture of 20 g of silicone elastomer SYLGARD 184 (Dow Corning) and 1 g of crosslinking agent, spin-coated at 1500 RPM for 60 sec, aiming to obtain a PDMS layer of 50 µm. The PDMS-coated FL-W was incubated at room temperature for 20 min, and baked at 80 °C for 18 minutes. The CL-W was put in a square Petri dish and covered with a degassed mixture of 40 g of silicone elastomer and 8 g of crosslinking agent, degassed for 30 min in a vacuum chamber and baked at 60 °C for 30 min. Patterned PDMS was cut with a blade and detached from the mold. Inlet ports were generated via a biopsy punch (1.0 mm OD, Harris Uni-Core). The CL was manually aligned with the FL, by visually overlapping the microchambers of the two layers. Bonding between the two layers was ensured by the presence of different amounts of crosslinking agent, which migrated from the CL to the FL. Next, metallic connectors (0.8/1.2 mm ID/OD, Phymep) were inserted into the inlet and outlet ports of the CL. To consolidate the assembly, the bonded FL and CL were further covered with a degassed mixture of 30 g of silicone elastomer and 3 g of cross-linking agent, further degassed in a vacuum chamber for 30 min, and baked at 80 °C overnight. PDMS was detached from the FL-W, metallic connectors were removed, and the inlet and outlet ports of the FL were punched (1.0 mm OD, Harris Uni-Core).

Next, a large coverslip (112 x 100 x 0.2 mm) was cleaned with isopropanol, dried, and bonded to the PDMS assembly by oxygen plasma treatment. Specifically, the coverslip was put inside a plasma chamber, the vacuum was created at 0.1 mbar, and oxygen was inflated at 0.25 mbar for 60 sec. Afterward, the PDMS assembly was put inside the plasma chamber with the microchannels upward. The same cycle was run for a duration of 30 sec. Just after plasma activation, the PDMS assembly was placed on the coverslip, covered with a metal weight (130 x 100 x 30 mm) of 500 g and incubated at 80 °C overnight to complete bonding.

To fix the metallic reservoirs (Extended Data Fig. 1g) on the microfluidic device, 5 g of silicone elastomer and 0.5 g of crosslinking agent were mixed and degassed and poured on the PDMS assembly using a 2-mL syringe, after closing the reservoir ports with metallic connectors. Next, the metallic reservoirs were placed on the surface of the device, and PDMS was crosslinked at room temperature for 24 h. Following this first incubation step, the metallic connectors were removed, and the half-kilo metal weight was placed on top of the assembly, which was further incubated at 80 °C overnight. Finally, the platform was decontaminated in a UVO-Cleaner (Jelight) for 20 min and stored in a sterile petri dish.

A pair of acrylic lids were manufactured, to seal each reservoir and inflate air, for injecting the contents of the reservoirs into the device microchambers. Each injector was made by gluing two 2-mm acrylic rectangles patterned with a groove and rimming thermoplastic polyurethane tape around the perimeter of the lid (Extended Data Fig. 1h). Finally, a metal connector, plugged into a 7-cm-long Tygon tubing with a female Luer lock at the opposite end, was also glued within the groove of the lid. The tubing was then connected to the third channel of the flow controller to drive injection. The metal and acrylic holders (Extended Data Figs. 1e and f), used to mount the platform on the microscope stage, were also manufactured.

### Characterization of injection from the reservoirs

Qualitative validation was carried out with food dyes. The platform was first manually filled with 7H9 medium from the outlet, until a drop of the medium was visible at the inlet ports of the reservoirs. Next, the reservoirs were alternately filled with red and blue food dyes and closed by screwing the injection lids. The inlet port of the FL was connected to the medium bottle that was closed with a pressure lid (Fluigent) and flangeless fittings (IDEX Health & Science), which were connected to a Tygon tubing (Saint-Gobain), in turn connected to a metallic fitting on the other end. The outlet port was connected to an empty waste bottle in the same way. The inlet port of the CL was connected to a 50-cm Tygon tubing via a metallic connector, and the other end of the tubing was connected to the flangeless fitting of the water reservoir. The outlet port of the CL was connected to a 5-cm Tygon via a metallic fitting and locked with a plier. The platform was controlled from the multi-channel Microfluidic Flow Control System (MFCS, Fluigent), via the MAESFLOW Control software (Fluigent): channel (C) 1 drove the FL inlet; C2 drove the CL inlet; C3 drove the reservoirs injecting lids; and C4 drove the FL outlet. First the FL was perfused with 7H9 medium by setting C1 at 200 mbar and closing the outlet with a plier for 20 min to degas the FL fluidic network. Next, C1 was set at 100 mbar, C2 and C3 were set at 30 mbar, and C4 at 20 mbar for 10 min. To actuate the reservoirs, C1 was set at 60 mbar and C3 at 100 mbar for 30 min, after which the device was photographed.

Quantitative validation was carried out with fluorescein isothiocyanate (FITC). The device was pre-filled with 7H9 medium and connected as described above, and the reservoirs were alternately filled with a FITC solution (100 µM) and with non-fluorescent 7H9 medium. The platform was mounted on the microscope stage via the custom-made holders (Extended Data Figs. 1e and f). The FL was perfused with 7H9 medium by setting C1 at 200 mbar for 20 min and closing the outlet with a plier for degassing. Next, C1 was set at 100 mbar, C2 and C3 were set at 30 mbar, and C4 at 20 mbar. Pressures were further adjusted by ±15 mbar, aiming to achieve a flow rate of 160 µL/h at the FL inlet port and 150 µL/h at the FL outlet port. After 6 h, C1 was set at 60 mbar and C3 was set at 100 mbar, to start injection from the reservoirs for 6 h. To stop injection from the reservoirs and restore the main flow from the bottle of medium, C1 was reverted to 100 mbar, C2 to 35 mbar and C3 to 30 mbar. At the end of the injection sequence, the whole device was manually filled with 100-µM FITC solution, to image the fluorescence of undiluted dye. To determine the dilution that the solutions stored in the reservoirs undergo during injection, via mixing with the medium in the flow layer, fluorescence signals were measured in the channel section just preceding the microchambers during injection and compared to the signal obtained from undiluted FITC solution. We calculated a dilution of approximately 2.5 times. Fluorescence images were acquired on a DeltaVision Elite Microscope, equipped with a Plan-Apochromat 10x/0.45 M27 objective (Zeiss), using the FITC channel (Ex 475/28, Em 525/48 nm) at 50%T, for 100 msec.

### Dynamic microfluidic screening setup and imaging

All connecting fluidic material was autoclaved before use, and device preparation was carried out under biosafety cabinet. The inlet port of the FL was connected to 50-cm Tygon tubing via metallic fitting. The other side of the tubing was attached to the 7H9 medium bottle as described previously, also adding a flowmeter M size (Fluigent) in between. The inlet port of the CL was connected to a 50-cm Tygon tubing via metallic fitting, and the other end of the tubing was attached to the water reservoir as described previously. The outlet port of the CL was connected to a 5-cm Tygon via metallic fitting and closed with a plier. The outlet port of the FL was first used to manually seed the bacterial single-cell suspension, obtained from filtration through a 5-μm filter (Merck), and then to drain spent medium into the waste bottle, filled up for one fifth with a 2.6% sodium hypochlorite solution. About 400 µL were gently injected from a 1-mL syringe connected to a 5-cm Tygon tubing (1.2 mm ID, Saint-Gobain) via a female Luer-to-Barb fitting, and the opposite side of the tubing was connected to the FL outlet via metallic fitting. Following 10-min incubation at room temperature, the FL outlet was connected to a 50-cm Tygon tubing via metallic fitting. A flowmeter was connected to this outlet tubing, and the other end was connected to a 5-cm Tygon tubing with a flangeless fitting plugged into a waste bottle as described previously. Next, the reservoirs were filled with 300 µL of compounds to be screened, at a concentration 2.5 times higher than the final concentration to be tested, due to the dilution occurring with 7H9 medium during the injection phase. Compounds with a MIC > 500 μM were tested at a final concentration of 250 μM; compounds with a MIC ≤ 500 μM were used at a final concentration equal to 0.5X-MIC. DMSO was used as a negative control; mitomycin C (300 nM) was used as a positive control. To control injection, the two conditions at the extremities of the reservoirs were filled with FITC solution (100 µM). After filling, the reservoirs were sealed by screwing on the injection lids.

Finally, the loaded platform was mounted on the microscope stage via the custom-made holders and controlled via the MFCS (Fluigent) as described above: C1 drove the FL inlet; C2 drove the CL inlet; C3 drove the reservoirs injecting lids; and C4 drove the FL outlet. The FL was first perfused with 7H9 medium at 200 mbar for 20 min, and the FL outlet was closed with a plier for another 20 min for degassing the microfluidic network. Next, the plier was removed and C1 was set at 100 mbar, C2 and C3 at 30 mbar, and C4 at 20 mbar, with further adjustments of ±15 mbar, targeting a flow rate of 160 µL/h at the FL inlet port and 150 µL/h at the FL outlet port. After the pre-growth stage, compounds were injected by setting C1 at 60 mbar and C3 at 100 mbar for 6 h, and injection was stopped by reverting C1 to 100 mbar, C2 to 35 mbar, and C3 to 30 mbar.

Time-resolved microscopy was carried out on an inverted DeltaVision Elite Microscope (Leica), equipped with a UPLFLN100XO2/PH3/1.30 oil objective (Olympus). Exposure conditions: phase contrast 100% T, 150 msec; FITC (Ex 475/28, Em 525/48 nm) 50% T, 150 msec; mCherry (Ex 575/25, Em 625/45 nm) 50% T, 150 msec. Images were acquired at either 20- or 30-min intervals. In each microchamber between 5 and 10 fields of view were recorded, each one containing between 1 and 10 individual bacteria. Each experiment was performed at least twice independently. At the end of each experiment, the platform was decontaminated with 2.6% sodium hypochlorite and discarded; the two flowmeters were washed with 2.6% sodium hypochlorite, followed by 70% ethanol and sterile water for 1 hour; tubing, fittings and connectors were autoclaved.

### Time-resolved image analysis and effect size estimation

Movies derived from the dynamic microfluidic screening were analyzed at the microcolony level for all compounds and at the single-cell level for shortlisted compounds.

Data extraction for microcolonies: raw image stacks were separated by channel and smoothed, applying a gaussian convolution kernel. Whole-microcolony binary masks were created based on the background of red channel images, using K-means clustering to separate the regions of interest (ROI), containing cells, from the remaining background image. All objects in the image that were smaller than a single cell, detached from the main ROI, or located on the edge of the image were excluded from the final mask. Further, a shape complexity index (*perimeter/√Area*) was calculated for each ROI. To do so, a Sobel operator was applied to identify boundaries between microcolony and background. Only images with a shape complexity index higher than 14 (determined based on a subset of images) were kept. This process was repeated iteratively for each image in the stack (movie), and the final binary masks were visually inspected. Images were retained with an overall success rate of 54%. Growth rate of microcolonies was estimated based on size increase over time using a robust regression, assuming growth was exponential. Pixel intensities within the ROI were centered based on the average value of the image background and log_10_ transformed to reduce the effects of background noise. ROI were also used to estimate mean (µ_RecA-GFP_) and variance (σ^2^_RecA-GFP_) of fluorescence intensity. The latter was used as a final measure of cell-to-cell variation. The average of each index was computed in every colony across the four last time points of each stage, the pre-exposure and the exposure stage. Each microcolony dataset included: three variables of interest (fluorescence mean; fluorescence variance; and growth rate); two experimental factors (compound and exposure stage, pre- or during exposure); and two technical factors (experiment replicate and microchamber coordinates).

Data extraction for single cells: *M. smegmatis* single-cell analysis was carried on ImageJ (2.0.0-rc-59/1.51n)^56^, using ROI Manager macro at given time points. Individual cells were manually segmented using the selection brush tool, and background mean fluorescence intensity was subtracted from each ROI to obtain final single-cell µ_RecA-GFP_ intensities. As done for microcolonies, average and variance of fluorescence intensity were also calculated for single cells. Finally, each single-cell dataset included: two variables (fluorescence mean and fluorescence variance); two experimental factors (compound and exposure stage); and two technical factors (experimental replicate and microchamber coordinates).

Statistical analysis: Mixed-effects models were used to assess the effect of exposure to PTC. The experimental factors (compound and exposure stage) were set as fixed effects, whereas technical factors were set as random effects. Standardized effect sizes were calculated to assess the impact of exposure to PTC as compared to the negative control group (DMSO). Statistical significance of these effects was tested based on *post-hoc* comparisons. Large positive or negative effect sizes reflect strong increase or decrease, respectively of values upon PTC exposure as compared to DMSO. Data extraction and analysis were performed with the R software 4.0.2 (https://www.r-project.org/). Detailed functions and scripts used in this study were packaged and are available at https://gitlab.pasteur.fr/svolant/ptc-screening.

### In silico docking

Docking poses of M06 or its amino derivative, INH and MOX were generated on SeamDock docking web server^57^ (http://doi.org/10.5281/zenodo.4506970). The structure PDB 4BGF of the arylamine N-acetyltransferase from *M. tuberculosis* cross-seeded with the crystals of *Mycobacterium marinum* protein^30^, and the structure PDB 5BS8 of the DNA gyrase from *M. tuberculosis*^24^ were used to predict ligand binding. Either Vina or Autodock were used to assess protein-ligand interactions with following settings: grid spacing between 0.2 and 1Å; energy range at 5 kcal/mol; and exhaustiveness at 32.

### MIC evaluation by resazurin assay

Secondary cultures were grown up to OD_600nm_ 0.6-0.8, diluted to OD_600nm_ 0.005 using pre-warmed Middlebrook 7H9 broth, and 100 μL were dispensed into a 96- well plate, including negative and positive controls. Wells containing the highest drug concentration to be tested were filled with 200 µL of cell suspension and used for sequential two-fold dilutions. Plates were incubated under static conditions for 7 days at 37 °C, after which 10 µL of resazurin (0.01%) were added to each well. Plates were further incubated for 24 h and then visually inspected, considering blue as cidality and pink as viability. MIC was regarded as the lowest concentration of drug causing cidality.

### Checkerboard Assay

The efficacy of dual drug-combinations was tested by checkerboard titration methods using 96-well plates, as described for the MIC assay. The first top-left well was used for the negative control. The rest of the first row and the first column were used to dilute each compound alone, whereas the highest concentrations of the two compounds to be tested (A and B) were alternately placed in the second row and in the second column, to test crossed dilutions. The drug concentrations tested ranged from 30 to 0.03 mM, except for M04, which ranged from 300 to 0.3 µM. The Fractional Inhibitory Concentration (FIC) index for combinations of two compounds A and B was calculated according to the following equation: FIC index = (A/MIC_A_) + (B/MIC_B_), where A and B are the MIC of compound A and compound B in combination, and MIC_A_ and MIC_B_ are the MIC of compound A and compound B alone. The FIC index was interpreted as follows: synergy (≤ 0.5), additivity (0.5 < FIC ≤ 1), and indifference (> 1).

### Kinetic assay of drug-mediated mortality

Secondary cultures of *M. tuberculosis* H37Rv were grown up to OD_600nm_ 0.5 and diluted to OD_600nm_ 0.05 in fresh 7H9 medium containing 0.01% Tyloxapol. Drugs were added as follows: INH (MIC: 50 ng/mL, 2X MIC); RIF (156 ng/mL, 2X MIC); MOX (100 ng/mL, 2X MIC); EMB (5 µg/mL, 2X MIC); M06 (1.25 µg/mL, 1X MIC); or their combinations. Untreated and drug-treated cultures were re-incubated at 37 °C at 50 RPM and aliquots were withdrawn sequentially for plating different dilutions on 7H11 agar medium enriched with 10% OADC and 0.5% glycerol. CFU were quantified after 4 weeks of incubation at 37 °C.

### Toxicity assay

Confluent Vero cells were dissociated by trypsinization, washed, resuspended in complete DMEM medium at a concentration of 2.5*10^5^ cells/mL, and 100 µL were dispensed into a 96-well plate and incubated for 24 h at 37 °C in humidified 5% CO_2_ atmosphere. Each compound (except for A1, A2, A3, A5, C2 and D10) was tested at a maximum concentration of 8 mM or 0.5 mM, depending on reagent availability, and serially diluted. Cells were exposed to the compounds for 24 and 48 h. Survival was assessed by ATP luminescent cell viability assay (CellTiter-Glo®, Promega). Half maximal inhibitory concentration (IC50) was estimated by nonlinear regression analysis of compound concentration versus normalized response using GraphPad Prism 9.

### Isolation of PTC-resistant mutants

Secondary cultures of *M. tuberculosis* H37Rv were grown up to OD_600nm_ 1 and about 10^10^ cells were harvested and spread on 7H10 agar plates containing drug concentrations ranging from 4- to 20-fold the MIC. Plates were incubated at 37 °C for 4 weeks. The resistance phenotype was validated by assessing the MIC at least twice for isolated colonies.

### Genomic DNA isolation and whole-genome sequencing

PTC-resistant mutants were reinoculated in the presence of the compound (8X MIC) and 10 mL of exponentially growing cultures were used for genomic DNA isolation, as follows. Each cell pellet was heat-inactivated at 80 °C for 1 h. Inactivated cells were washed and resuspended in 450 µL of lysis solution (25 mM Tris-HCl pH 7.9; 10 mM EDTA 50 mM glucose), also adding 50 µL of lysozyme (10 mg/mL) and incubated at 37 °C overnight. Next, 100 µL of sodium dodecyl sulfate (10 %) and 50 µL of proteinase K (10 mg/mL) were added, mixed gently by inversion, and incubated at 55 °C for 30 min. Next, 200 µL of sodium chloride (5 M) were added and mixed gently by inversion, followed by 160 µL of preheated cetrimide saline solution (4 %), mixed and incubated at 65°C for 10 min. Genomic DNA was extracted twice, adding equal volume (1 mL) of chloroform: isoamyl alcohol (24:1), mixing by inversion, and centrifuging for 5 min at 13500 *g*. The aqueous layer was transferred to a clean tube (800 µL) and 560 µL (0.7 volumes) of isopropanol were added at room temperature, mixed by inversion until DNA precipitation. DNA was collected by centrifugation for 10 min and the pellet was washed with 70 % ethanol, dried and resuspended in 100 µL of TE buffer. Libraries for whole-genome sequencing (WGS) were built using a TruSeq DNA PCR-Free Low Throughput Library Prep Kit (Illumina, USA) following the manufacturer’s protocol. Quality control was performed on Agilent Bioanalyzer. DNA sequencing was performed on the Illumina MiSeq 250 platform using paired-end 150-bp reads.

### Variant calling

Variant calling analysis was performed with Sequana 0.9.8^58^, using the variant calling pipeline 0.10.0 built on top of Snakemake 6.1.1^59^. Briefly, whole genome sequencing reads were aligned to the reference genome of *M. tuberculosis* H37Rv v2 using bwa 0.7.17^60^. Sambamba 0.8.0^61^ was used to mark and filter duplicated reads. Variants were identified using Freebayes 1.3.2^62^ and annotated using SNPeff 5.0^63^. High quality single nucleotide polymorphisms (SNP) were selected using the following criterions: freebayesscore: 20; frequency: 0.7; mindepth: 10; forwarddepth: 3, reversedepth: 3, strandratio: 0.2. Quality control statistics were summarized using multiQC 1.10.1^64^.

### Total RNA extraction and sequencing

Secondary *M. tuberculosis* Erdman cultures were grown in Middlebrook 7H9 broth until exponential growth phase (OD_600nm_ 0.6) and were subjected to different compounds at a concentration of 10-fold the MIC for 6 h, in triplicate. DMSO was used as a negative control. For total RNA isolation, 10 mL of each culture were collected at 4200 x *g* for 15 min at 4 °C and frozen at −80 °C for 24 h. RNA was extracted by hot-phenol method and precipitated with 0.5 M LiCl and three volumes of cold 100% ethanol for 2 h at -20 °C. Samples were centrifuged at 13500 x *g* for 35 min at 4 °C and RNA was washed with 80% ethanol. RNA samples were air-dried for 30 min and resuspended in 20 µL of nuclease-free water. DNA was depleted by double DNase treatment, using the TURBO DNA-free kit (Ambion). The quality and integrity of RNA samples were checked on Agilent 2100 Bioanalyzer, and the amount was quantified on the Qubit fluorimeter (Thermo Fisher). Ribosomal RNA was depleted using the QIAseq FastSelect 5S/16S/23S kit for bacteria (Qiagen), according to manufacturer’s recommendations. Libraries were prepared using the TruSeq Stranded mRNA Library Preparation kit (Illumina) according to manufacturer’s protocol: (https://support.illumina.com/content/dam/illumina-support/documents/documentation/chemistry_documentation/samplepreps_truseq/truseq-stranded-mrna-workflow/truseq-stranded-mrna-workflow-checklist-1000000040600-00.pdf). The quality, integrity and amount of the libraries were checked with the Agilent 2100 Bioanalyzer and Qubit fluorimeter. Lastly, single-end sequencing was carried out on a NextSeq500 mid-output system (Illumina), 130 M reads.

### RNA-seq

RNA-seq datasets were produced from three independent experimental replicates. Sequana 0.9.8 rnaseq pipeline 0.10.0 built on top of Snakemake 5.8.1 was used to perform data analysis. In short, reads were trimmed from adapters using Cutadapt 2.10^65^, then mapped to *M. tuberculosis* Erdman genome (ASM35020v1) using bowtie 2.2.2^66^. FeatureCounts 2.0.0^67^ was used to produce the count matrix, assigning reads to features with strand-specificity information. Quality control statistics were summarized using MultiQC 1.8. Statistical analysis on the count matrix was performed to identify DEGs, comparing *M. tuberculosis* treated with a given drug relative to the DMSO control condition. Clustering of transcriptomic profiles were assessed using a Principal Component Analysis. Differential expression testing was conducted using DESeq2 library 1.24.0^68^ with SARTools 1.7.0^69^, indicating the significance (Benjamini-Hochberg adjusted p-values, false positive rate < 0.05) and the effect size (fold-change) for each comparison. GO enrichment analysis was carried out using the Gene Ontology Resource powered by PANTHER (http://geneontology.org/)^70, 71^.

### Real-time quantitative PCR

To validate transcriptional silencing of *nat* by CRISPRi, secondary cultures of GMT36 and related control were grown up to OD_600nm_ 0.6-0.8, diluted to OD600 0.05 and induced with 100 ng/mL of anhydrotetracycline (ATC) for 4 days in duplicate. To quantify the expression of genes implicated in stress response, *M. tuberculosis* H37Rv and NAT mutants (S9, S10 and S11) were cultured in Middlebrook 7H9 broth until OD_600_ 0.6, in duplicate. Next, cultures were exposed to M06 (10X MIC) for 6 h. Total RNA was extracted by hot-phenol method and DNA depleted, as described above. cDNA was generated from 150-200 ng of total RNA using SuperScrip IV (Invitrogen) and random hexamers (Thermo Fisher), according to manufacturer’s instruction. qRT-PCR analysis was carried out using the SYBR Green PCR Master Mix (Applied Biosystems), with 0.3 μM primers, and 1 μL cDNA diluted 1:4. Absolute quantification was run on the LightCycler®480 Instrument (Roche Life Science): activation at 50 °C for 2 min (ramp rate °C/s 4.8) followed by 95 °C for 10 min; 40 amplification steps at 95 °C for 15 sec (ramp rate °C/s 4.8), followed by 60 °C for 30 sec (ramp rate °C/s 2.5) and 72 °C for 30 sec (ramp rate °C/s 4.8); melting curve at 60 °C for 15 sec (ramp rate °C/s 2.5), followed by 95 °C for 15 sec (ramp rate °C/s 0.29). Genomic DNA of *M. tuberculosis* was serially diluted to generate standard curves and calculate transcript copy numbers.

### Snapshot microscopy and single-cell image analysis

Secondary cultures of *M. tuberculosis* H37Rv, NAT mutants (S9, S10 and S11) and GMT35 NAT-overexpressing strain were cultured in Middlebrook 7H9 broth until OD_600_ 0.5, in duplicate. Cultures were treated for 24 h with DMSO (CT), H_2_O_2_ (30 mM), or with different drugs at 10X-MIC concentration: M06 (12.5 µg/mL); MOX (500 ng/mL); INH (250 ng/mL); rifampicin (RIF) (800 ng/mL). Next, bacilli were stained with either CellROX (5 µM) or FM464 (5 µg/mL) dyes for 30 min and imaged by phase-contrast and fluorescence using an inverted DeltaVision Elite Microscope (Leica) equipped with an UPLFLN100XO2/PH3/1.30 objective (Olympus). All samples were prepared dispensing 0.6 μL of bacterial suspension between two #1.5 coverslips, sealed with glue. Exposure conditions: phase-contrast 50% T, 100 msec; Cy5 (Ex 632/22, Em 679/34) 50% T, 100 msec for CellROX detection; TRITC (Ex 542/27, Em 597/45) 50% T, 100 msec for FM464 detection. Individual bacilli were automatically segmented in phase contrast, using Omnipose^37^ for cell segmentation in combination with a custom-trained model. The latter was obtained from training on a dataset consisting of the ground truth dataset provided as part of Omnipose and additional manually labeled images of *M. tuberculosis*. A Python notebook was written to extract single-cell fluorescence and cell area from snapshot images using the Omnipose-derived segmentation masks. Inverted masks were used to extract background fluorescence, which was subtracted from the fluorescence of each cell.

### Liquid chromatography and mass spectrometry (LC-MS) analysis of M06 metabolites

Secondary cultures of *M. tuberculosis* H37Rv were grown up to OD_600nm_ 0.8 in duplicate and treated either with 18.75 µg/mL *of M06* (15X MIC) or with corresponding volume of DMSO as negative control. At each time point (0, 24, and 48 hours), 20 mL of culture were collected at 4200 *g* for 10 min at 4 °C. Supernatants were filtered through 0.22-µm filtration units and stored at 4 °C. M06 and its metabolites were extracted from 20 mL of culture supernatant three times, by adding same volume of water and same volume of ethyl acetate, which was then evaporated under vacuum and the residue was dissolved in 100 µL of acetonitrile. LC/MS analysis was performed using a Q Exactive mass spectrometer (Thermo Scientific). The instrument was equipped with an electrospray ionization source (H-ESI II Probe) coupled to an Ultimate 3000 RS HPLC (Thermo Scientific). A Thermo Scientific Hypersil GOLD aQ chromatography column (100 mm x 2.1 mm; 1.9 µm particle size) was prewarmed at 30 °C and injected with sample. The flow rate was set to 0.3 mL/min and the column was run for 3 min in isocratic eluent consisting of 5% acetonitrile in water containing 0.1% formic acid; 5 to 100% from 3 to 8 minutes. Full MS in positive mode was used with resolution set at 70000, max IT = 240 ms and AGC Target = 1.10^6^.

### Thin layer chromatography of lipids, fatty and mycolic acids

Secondary cultures of *M. tuberculosis* H37Rv, S10 NAT-mutant, and GMT35 NAT-overexpressing strain were cultured in Middlebrook 7H9 broth supplemented with 10% albumin-dextrose-catalase and 0.05% Tween 80, shaking at 120 RPM, until OD_600_ 0.15-0.2. Cultures were left untreated or treated with M06 (MIC: 1.25 µg/mL), MOX (50 ng/mL), or INH (MIC: 25 ng/mL), at different concentrations relative to the MIC: 1X; 5X; 10X; 25X; or 50X the MIC. At the same time cultures were labelled with either 0.5 or 1 µCi/mL of ^14^C-acetate (ARC; specific activity 106 mCi/mmol) and incubated at 37 °C under static conditions for 24 h. Lipids and mycolic acids were also analyzed in wild type, NAT-mutant and NAT-overexpressing strains, metabolically labeled with 1 µCi/mL of ^14^C-acetate as above, during early (OD_600_ 0.15) and mid (OD_600_ 0.5) exponential phase.

Lipids were extracted from 100 µl culture aliquots with 1.5 mL chloroform : methanol (2 : 1) at 65 °C for 3 h. After extraction, 150 µL of ddWater was added, the samples were mixed, centrifuged at 3 000 x *g* for 3 min and lower organic phase was removed and dried under nitrogen. Extracted lipids were washed with chloroform : methanol : ddWater (4 : 2 : 1), dried with nitrogen, resuspended in 50 µL chloroform : methanol (2 : 1) and 5 µL were loaded on silica gel 60 F254 plates (Merck). To separate trehalose esters of mycolates (TDM; TMM) and phospholipids (PE; CL), a mixture of chloroform : methanol : ddWater (20 : 4 : 0.5) was used as the eluent. To separate TDM, TMM and phosphatidylinositol mannosides (PIMs), a mixture of chloroform : methanol : NH_4_OH : ddWater (65 : 25 : 0.5 : 4) was used as the eluent. ^14^C-labeled lipid profiles were visualized using an AmershamTM TyphoonTM Biomolecular Imager.

For mycolic acids extraction, 100 µL cultures were treated with 1 mL of tetrabuthylammonium hydroxide (15%) and incubated at 100 °C overnight. Samples were methylated by adding 1.5 mL of dichloromethane, 1 mL of ddWater, and 150 µL of iodomethane, and incubated at room temperature under rotation for 4 h. Methylated samples were washed twice with ddWater, and organic phases were dried with nitrogen. Fatty and mycolic acid methyl esters (FAME and MAME) were extracted with 2 mL of diethyl ether, dried, and resuspended in 50 µL chloroform : methanol (2 : 1) and 5 µL were loaded on TLC plate, which was developed in n-hexane : ethyl acetate (95 : 5; 3 runs). Unsaturated forms of MAME were analyzed on Ag-impregnated silica plates. ^14^C labeled FAME and MAME were visualized as described above. Cell-envelope components were quantified from TLC autoradiographs using ImageJ^56^ and normalized to untreated samples.

### Intracellular efficacy imaging assay

Confluent Raw 264.7 macrophages were diluted to a concentration of 5*10^4^ cells per mL in complete DMEM without phenol red and without antibiotics, and 70 µL were seeded into each well of an 18-well µ-Slide (ibidi), and incubated at 37 °C in humidified 5% CO_2_ atmosphere for 24 h. Macrophages were infected with our *M. tuberculosis* ribosomal reporter, also constitutively expressing a red marker (rRNA-GFP_DsRed2; GMT17)^32^. Secondary *M. tuberculosis* cultures were grown up to OD_600_ 0.4-0.6, diluted in complete DMEM without phenol red to OD_600_ 0.0025 (MOI 1:10), and medium was replaced with the bacterial cell suspension, except for uninfected control wells. After 4 hours of incubation at 37 °C in humidified 5% CO_2_ atmosphere, infected cells were washed five times with complete DMEM without phenol red, and drugs were added as follows: INH (MIC: 50 ng/mL, 2X MIC); RIF (156 ng/mL, 2X MIC); M06 (1.25 µg/mL, 1X MIC); or their combinations. DMSO was used in untreated control wells. Cells were re-incubated for 6 days and stained with DRAQ7 mortality dye (3 µM) before imaging. Six images were acquired for each field of view through a z-stack of 6 µm, starting the scan from the middle of the sample, using a 100X oil immersion objective, 1.4NA, WD 0.12mm (Olympus). Exposure conditions on the Delta Vision microscope: bright field 10% T, 100 msec; FITC (Ex 475/28, Em 525/48) 32% T, 50 msec; TRITC (Ex 542/27, Em 597/45) 32% T, 50 msec; Cy5 (Ex 632/22, Em 679/34) 32% T, 50 msec. Image stacks were projected by maximum-intensity method in SoftWorx, and processed using a customized ImageJ macro, which automatically segments intracellular bacterial foci based on their red fluorescence. Derived masks were used to measure size and rRNA-GFP fluorescence normalized to size (as a proxy for metabolic activity) of intracellular bacilli. Inverted masks were used to subtract the background green fluorescence from each ROI.

### Statistics

Unless otherwise specified, plots and statistical analyses were done with GraphPad Prism 9.5.0 (525). Plots merge datasets deriving from at least two independent replicates, and significant *P* values, sample sizes and statistical tests are reported in each legend.

## Data availability

We have no restriction on data availability. Source data are either available in the article or were made publicly available in different repositories. Time-resolved image stacks derived from PTC screening (Source Data 2) are available at DRYAD, under the unique digital object odentifier doi:10.5061/dryad.r4xgxd2j8. Raw data from whole transcriptome and whole genome sequencing are available at the EMBL’s European Bioinformatics Institute, under the following EBI accession numbers: E-MTAB-12306 (RNA-seq) and E-MTAB-12307 (WGS). All other requests for reagents should be addressed to and will be fulfilled by the corresponding author, upon reasonable request and completion of a Materials Transfer Agreement.

## Code availability

Detailed functions and scripts used for the analysis of PTC screening were packaged and are available at https://gitlab.pasteur.fr/svolant/ptc-screening/. The custom Omnipose model and Python notebook used for single-cell snapshot analysis are available at https://gitlab.pasteur.fr/iah-public/automated_segmentation_mycobacterium_tuberculosis_snapshots.

## Acknowledgements

We thank Guillaume Duménil (Pathogenesis of vascular infections, Institut Pasteur, Paris, France), Katarína Mikušová (Comenius University in Bratislava, Slovakia), and Pietro Slavich (LPTHE, Paris, France) for critical readings of the manuscript. We thank Samy Gobaa for providing us access to the Biomaterials & Microfluidics core facility of the Institut Pasteur. We thank the Biomics Platform, C2RT, Institut Pasteur, Paris, France, supported by France Génomique (ANR-10-INBS-09-09) and IBISA for assistance in RNA-Seq and WGS. The mutant strains in Supplementary Table 5 were obtained from the Collection of the Institut Pasteur (CIP), Biological Resource Center of Institut Pasteur (CRBIP), Paris, France. This work was supported by the following grants to G.M.: French Medical Research Foundation grant ING20160435202; French National Research Agency grant ANR-17-CE11-0007-01 PersisTB; French National Research Agency grant ANR-10-LABX-62-IBEID; and Institut Pasteur core funding. The work was further supported by the French National Research Agency grant ANR-21-CE15-0045 TREATABLE to G.M. and C.R., and by the Slovak Research and Development Agency grant n. APVV-19-0189 and the OPII, ACCORD, ITMS2014+: 313021X329, co-financed by ERDF to J.K.

## Author contributions

Conceptualization: G.M.; M.C. and S.R.L. Methodology: M.M.; P.C.; S.V.; G.M. Software: P.C.; S.V.; E.K.; M.A.; M.M.; L.P.; G.M. Validation: M.M.; M.C.; P.C.; S.V.; E.K.; N.G.; J.K.; S.V.G.; G.M. Formal analysis: M.M.; P.C.; S.V.; G.M.; N.G.; L.P.; J.K.; M.Z.; S.V.G.; M.C. Investigation: M.M.; M.C.; G.M.; N.G.; J.K.; M.Z.; S.V.G. Resources: M.M.; M.C.; G.M.; O.H.; C.R.; P.D.; C.L.; C.T.; A.L.; Y.A.; S.R.L.; J.K.; M.Z.; S.V.G.; F.B.; P.C.; S.V.; E.K.; M.A.; J.Y.T. Data curation: M.M.; P.C.; S.V.; E.K.; G.M.; M.C.; Y.A. Writing—original draft: G.M. Writing—review and editing: all authors. Visualization: G.M.; M.M.; P.C.; S.V.; E.K.; J.K.; S.V.G.; Y.A. Supervision: G.M.; C.R.; P.D.; C.L.; S.R.L.; J.K.; J.Y.T. Project administration: G.M. Funding acquisition: G.M.; C.R.; J.K.

## Competing interests

G.M. and M.M. are designated as inventors in the pending international patent application WO 2020/229629 filed by the Institut Pasteur. M.C., P.D., and C.R. are designated as inventors in the pending international patent application WO/2019/138084 filed by the Institut Pasteur, Université de Caen Normandie, and Université Felix Houphouët-Boigny.

## Extended Data Figures

**Extended Data Figure 1.**
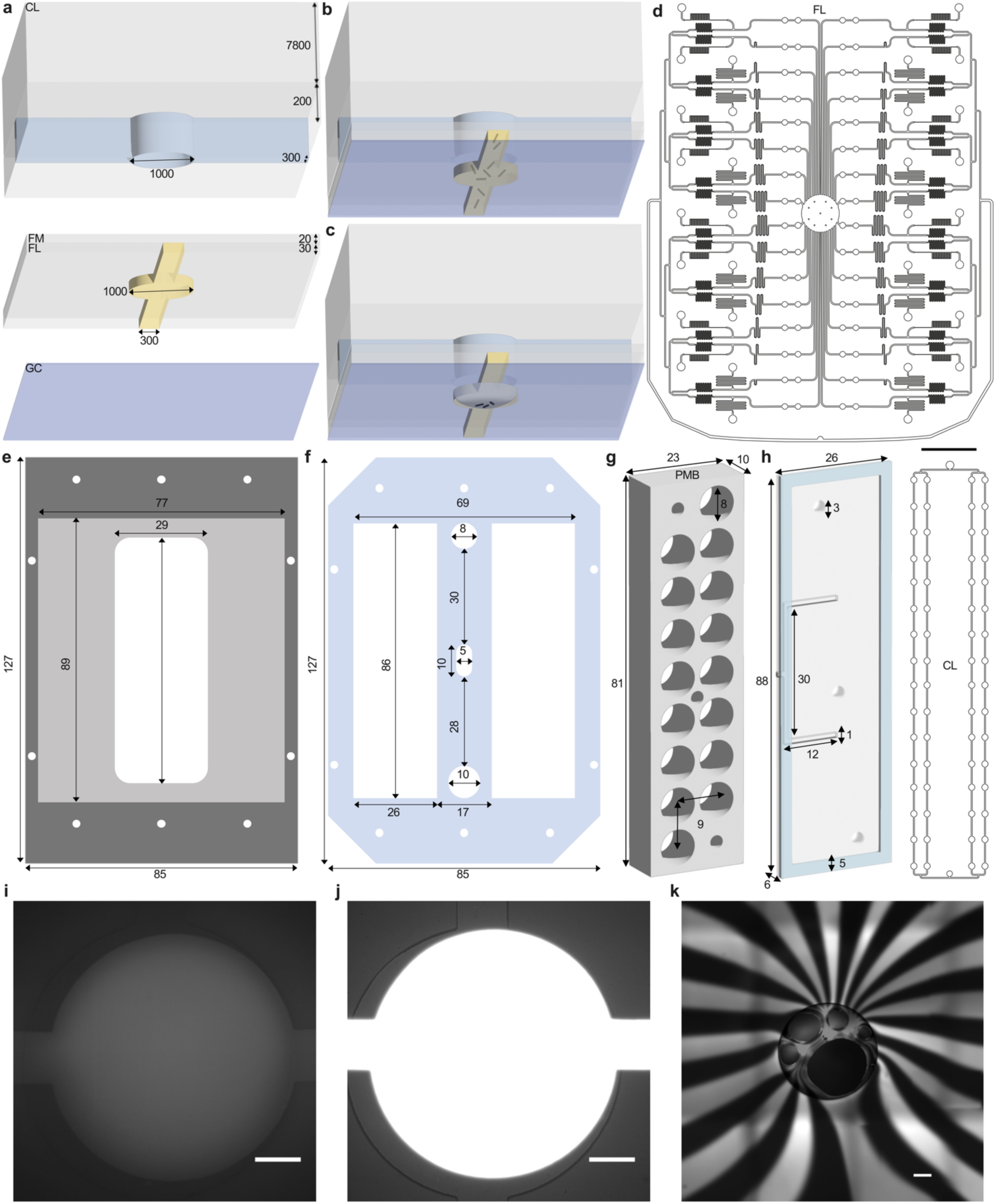
Design and structural components of the 32-condition microfluidic platform. **a**,**b**,**c** Not-to-scale axonometric projections of the components of the microchamber (viewed from the bottom). From top to bottom (**a**): control layer filled with water (CL, light blue); flexible membrane (FM); flow layer filled with growth medium (FL, yellow); glass coverslip (GC). Numbers represent dimensions in µm. Microchamber assembled layers, with bacteria (black rods) perfused in the FL, without (**b**), or with (**c**) pressure applied from the CL. Bacteria are trapped between FM and GC as the pressure increases in the CL. **d**, Mask design of the FL and CL. Scale bar, 10 mm. **e**,**f**, Schematics of the metal (**e**) and acrylic (**f**) holders used to mount the 32-condition device on the stage of an inverted wide-field microscope. The maximum area corresponds to a standard multi-well plate. Numbers indicate dimensions in mm. **g**,**h**, Axonometric projections of the perforated metal block (PMB) forming 32 reservoirs (**g**) and of its sealing lid, used for air injection (**h**). Numbers represent dimensions in mm. **i**,**j**,**k**, Representative fluorescence pictures of microchambers perfused either with non-fluorescent medium (**i**) or with a FITC solution (**j**), and of the outlet port of the device (circle) receiving non-fluorescent and fluorescent medium from the outlet channels (alternate dim and bright segments) of each pair of microchambers, showing proper separation of the conditions (**k**). Scale bars, 200 µm.

**Extended Data Figure 2.**
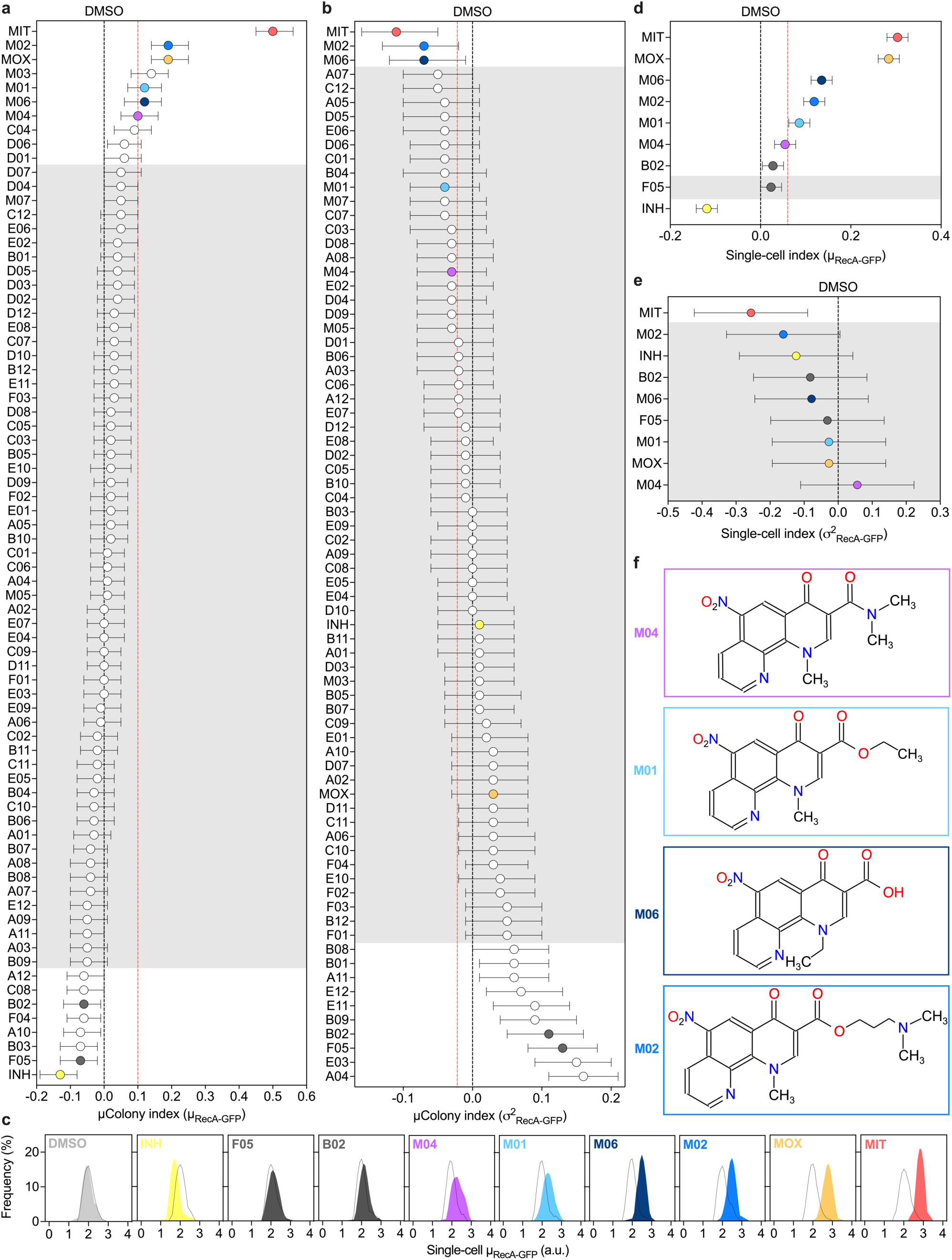
Effect-size estimates of the impact of PTC on *M. smegmatis* RecA-GFP. **a**,**b**,**d**,**e**, Comparison of the size of PTC effects on (**a**,**d**) average (µ_RecA-GFP_) and (**b**,**e**) variance (σ^2^_RecA-GFP_) of fluorescence levels as recorded in microcolonies (**a**,**b**) and in single-cell (**d**,**e**) setups. Effect sizes reflect difference between DUR versus PRE stages. Black dashed lines represent negative control DMSO. Red dashed lines represent 20% of the effect of MIT (red circle). Error bars represent mean ± confidence interval, from 2 to 15 independent experiments (5 < *n* < 86 microcolonies per condition; and 90 < *n* < 747 single cells per condition). White and gray areas represent significant and non-significant indices, respectively, which were estimated by mixed-effects models (Supplementary Table 2; Methods). **c**, Distribution of log_10_ single-cell µ_RecA-GFP_ at the end of PRE (white) and DUR (colored) stages (90 < *n* < 349 single cells, from at least two independent experiments). **f**, Chemical structure of PTC hits.

**Extended Data Figure 3.**
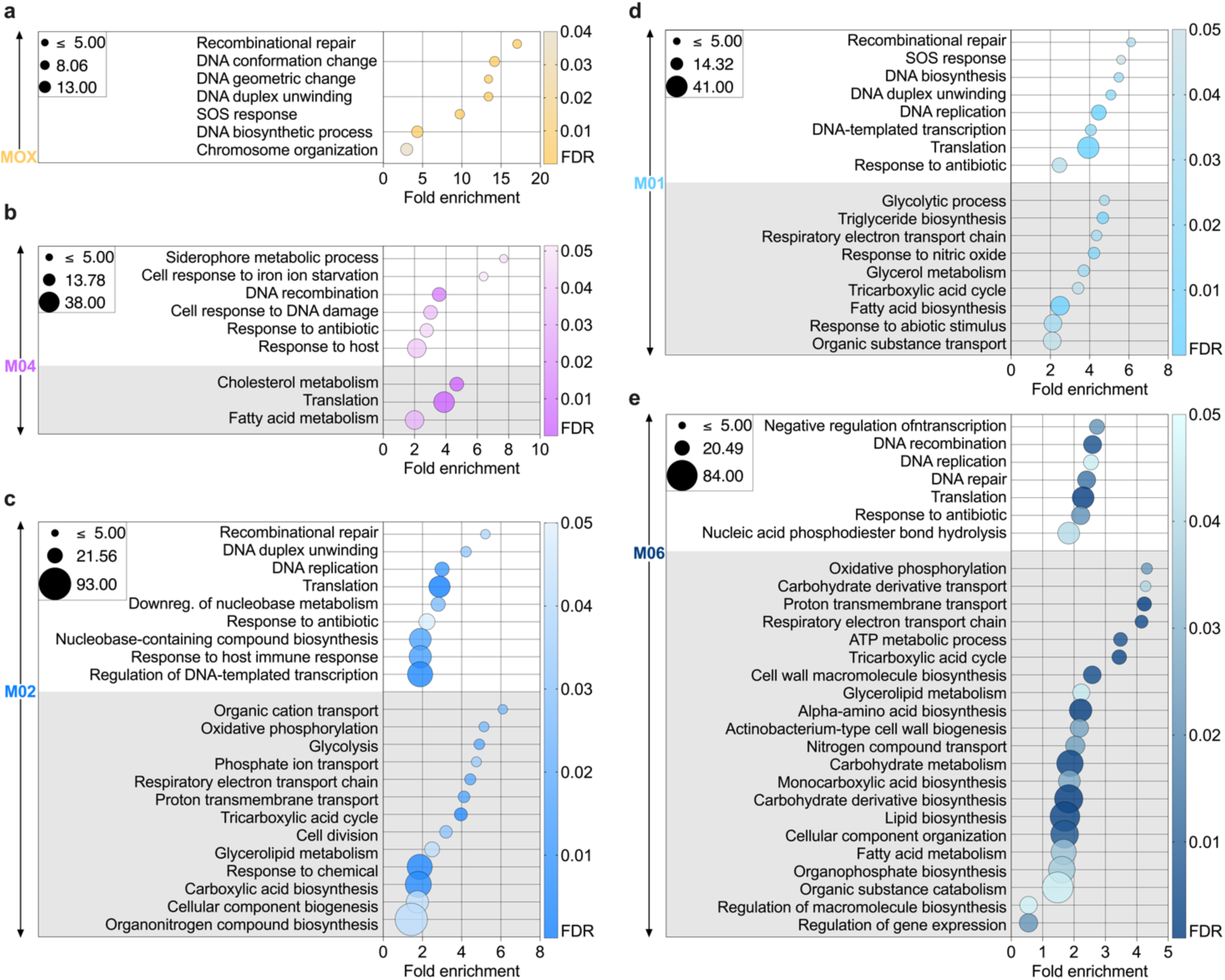
GO enrichment analysis of DEGs in *M. tuberculosis* treated with PTC hits and MOX. **a−e**, Complete GO biological processes were identified by overrepresentation test of significantly up-regulated genes (white shading) and down-regulated genes (gray shading), upon treatment with MOX (**a**); M04 (**b**); M02 (**c**); M01 (**d**); and M06 (**e**), against all *M. tuberculosis* genes in PANTHER database. Raw *P*-values were determined by Fisher’s exact test, and FDR < 0.05 was calculated by Benjamini-Hochberg procedure. Bubble plots reports the most specific functional categories sorted by fold enrichment. The entire hierarchies of significantly enriched functional categories for up- and down-regulated genes are also available (Supplementary Table 4).

**Extended Data Figure 4.**
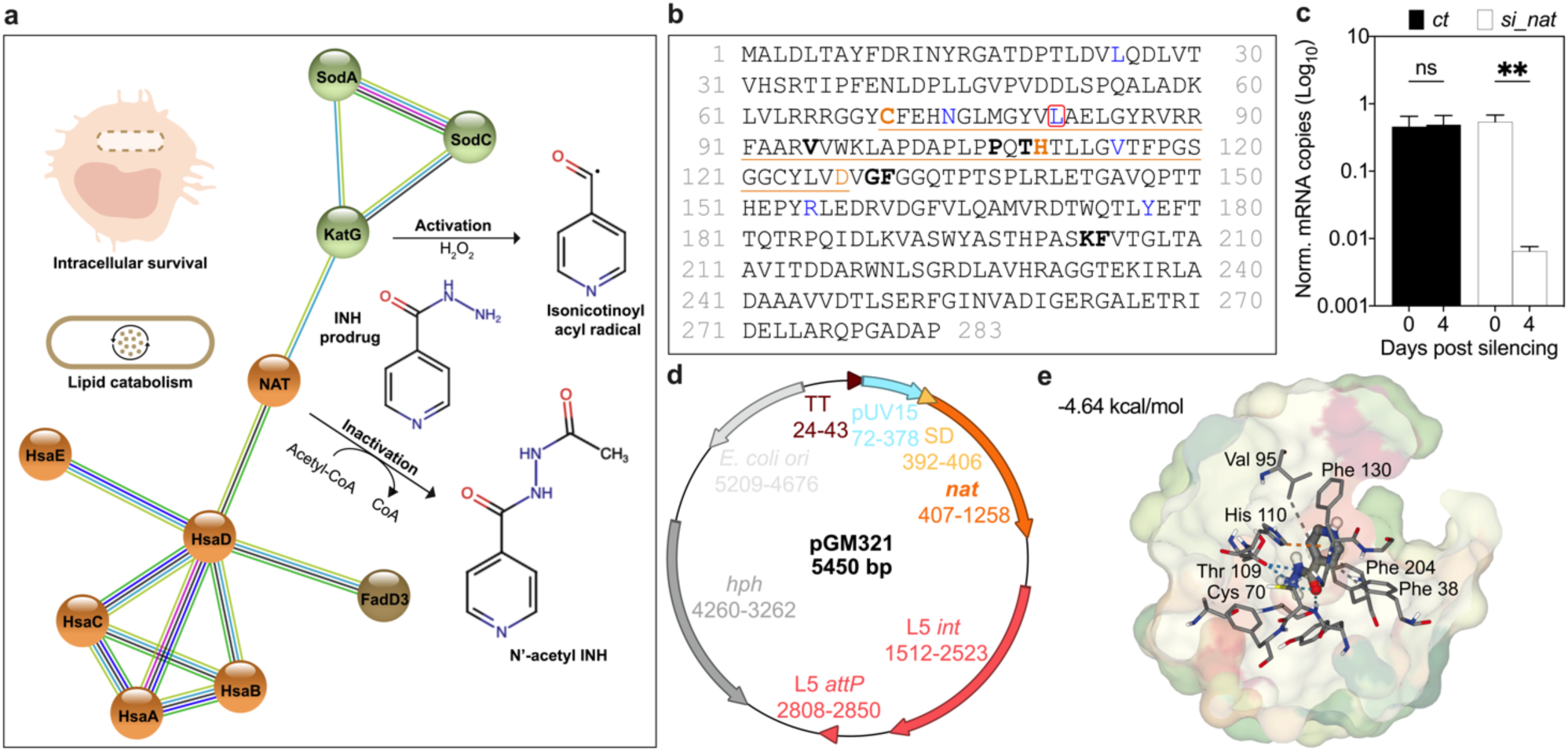
Role of NAT in mycobacteria. **a**, Highest confidence STRING interaction network of *M. tuberculosis* NAT, with no more than 5 interactors in the first shell and maximum seven interactors in the second shell (https://string-db.org/). Colored network edges indicate the type of interaction based on evidence: text mining (light green); experiments (pink); databases (light blue); co-expression (black); gene neighborhood (green); co-occurrence (blue). Activation and inactivation of INH prodrug by KatG and NAT, respectively, are sketched on the right of the network and other roles of NAT are sketched on the left. **b**, *M. tuberculosis* NAT protein sequence, with numbered amino acids. The active site is underlined, with residues belonging to the catalytic triad colored orange, and residues implicated in protein structure stability colored blue. Leucine 81, which is deleted in PTC-resistant mutants, is circled in red. Residues that potentially interact with M06 are bolded. **c**, qRT-PCR analysis of *M. tuberculosis nat* before and after induction of a dCas9 silencing system together with a scramble sgRNA (*ct*) or with an anti-*nat* sgRNA (*si_nat*). Transcripts are normalized to total RNA and *sigA* mRNA copies. Error bars represent mean ± SD (*n* = 2). Significance between 0- and 4-day post induction by two-way ANOVA followed by Šidák correction for multiple comparisons: \*\**P* = 0.0017. **d**, Map of the L5-phage integrative plasmid pGM321. A mycobacterial Shine Dalgarno (SD) and the *nat* open reading frame are cloned downstream of the strong pUV15 promoter. **e**, In-silico docking pose of CPK-colored INH in the catalytic pocket of NAT. Binding affinity is indicated (kcal/mol). Predicted amino acid residues in hydrophobic contact (dark gray); cation-pi interaction (orange); and linked by either weak (light gray) or strong (blue) hydrogen bonds with INH atoms are shown.

**Extended Data Figure 5.**
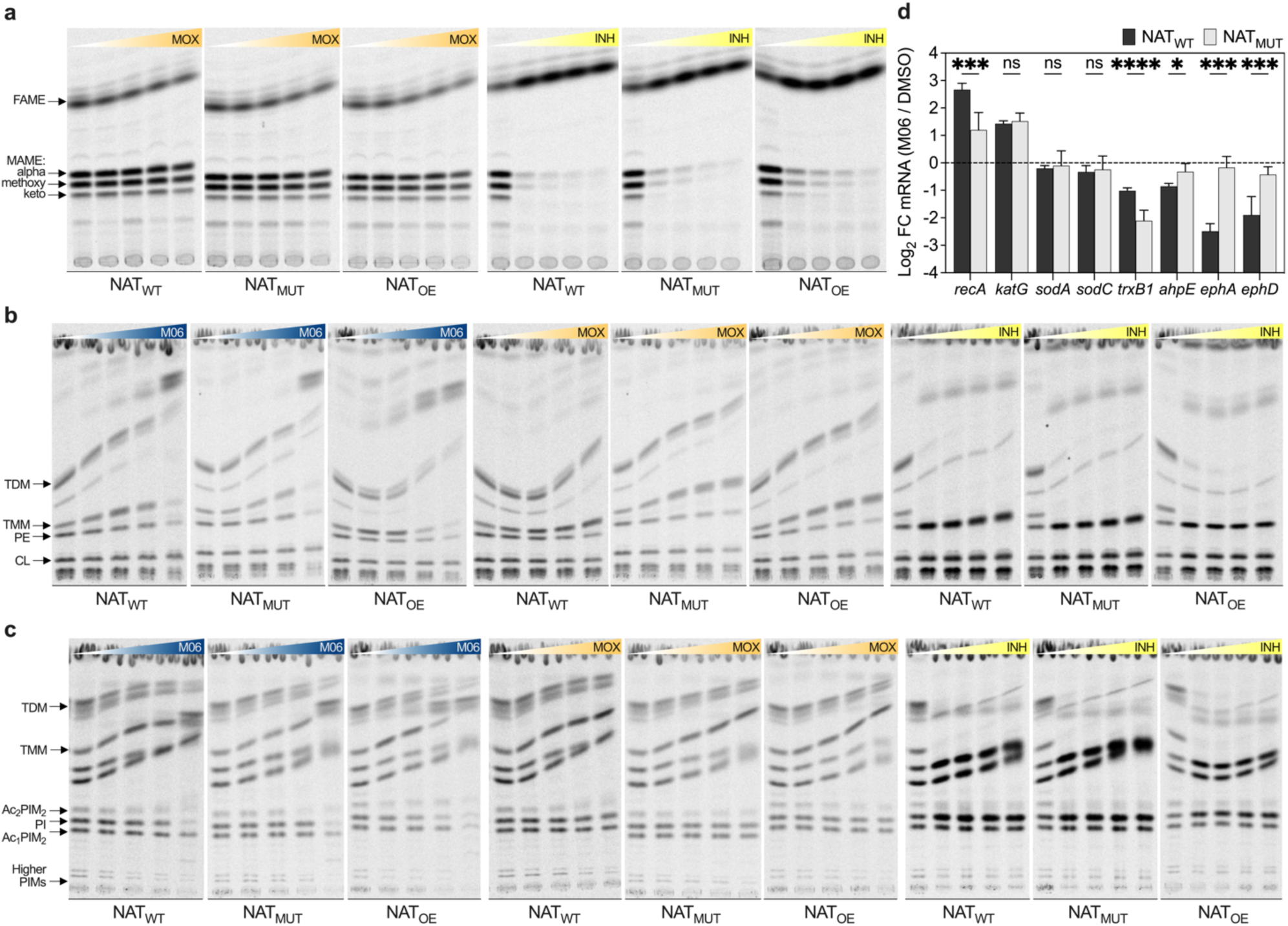
Lipid and transcriptional changes in drug-stressed *M. tuberculosis*. **a−c**, Representative TLC of fatty/mycolic acids methyl esters (FAME/MAME) and lipids from exponentially growing *M. tuberculosis* WT (NAT_WT_); S10 mutant (NAT_MUT_); and NAT overexpressing strain (NAT_OE_). Color gradients indicates from left to right untreated bacilli (white) or treatment with increasing concentrations of M06 (blue, MIC: 1.25 µg/mL), moxifloxacin (orange, MIC: 50 ng/mL) and isoniazid (yellow, MIC: 25 ng/mL) equal to 5X, 10X, 25X and 50X the MIC. FAME and different types of MAME (**a**). Trehalose dimycolate (TDM); trehalose monomycolate (TMM); phosphatidylethanolamine (PE); cardiolipin (CL) (**b**). TDM; TMM; phosphatidylinositol (PI); phosphatidylinositol mannosides (PIM); acylated forms of PIM (Ac_2/1_PIM_2_) (**c**). **d**, qRT-PCR analysis of *M. tuberculosis* WT (NAT_WT_) and averaged S9, S10 and S11 NAT mutants (NAT_MUT_). Transcripts are normalized to total RNA and *sigA* copies and are expressed as log_2_ FC between 24-hour treatment with M06 versus DMSO. Error bars represent mean ± SD (*n* = 2). Significance by multiple unpaired Welch *t*-tests, corrected by Holm-Šidák method: \**P* = 0.013; \*\*\**P* < 0.0005; \*\*\*\**P* = 0.000004.

**Extended Data Figure 6.**
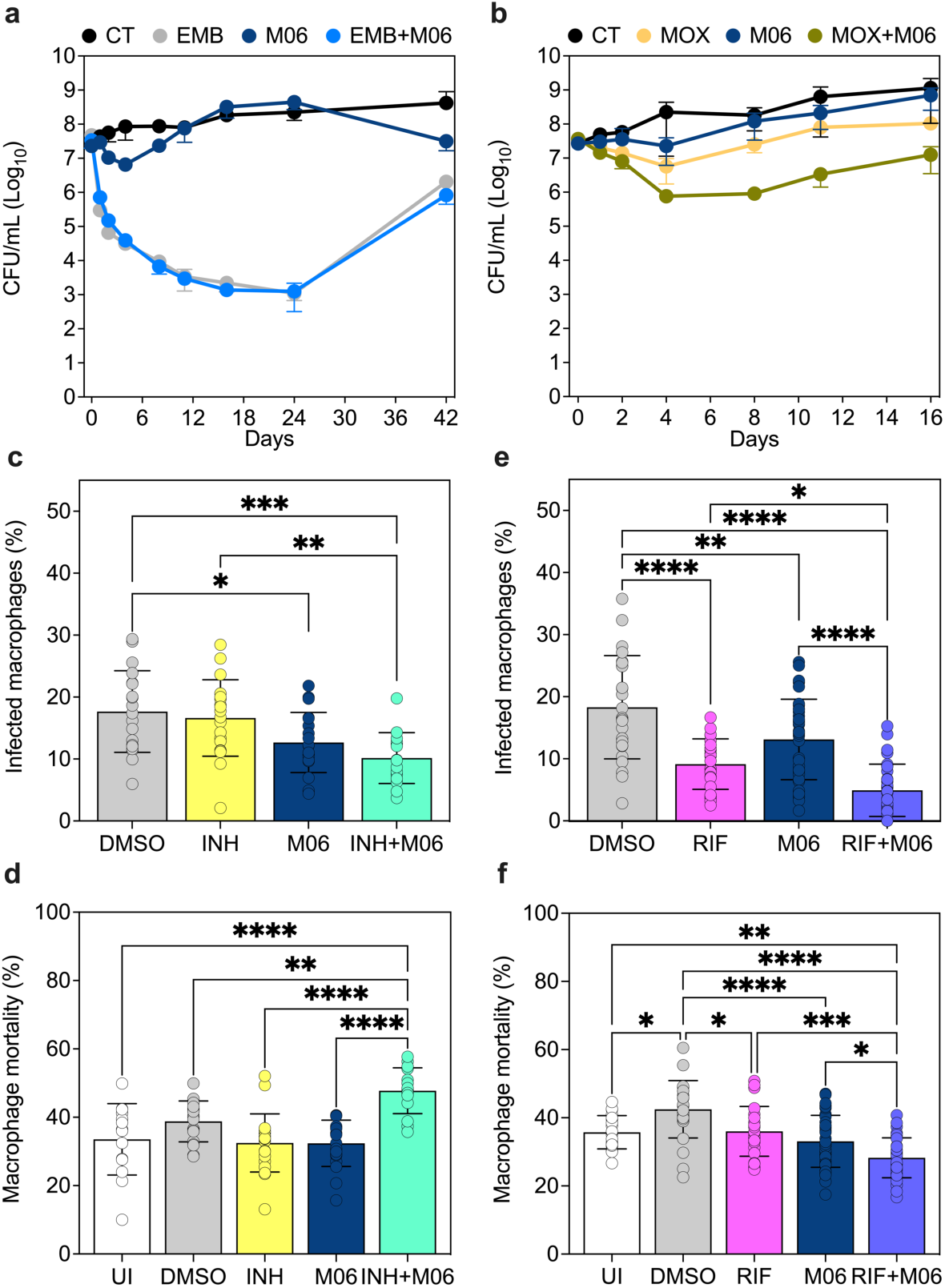
Analysis of M06 activity in combination. **a,b**, Drug activity in exponential-phase *M. tuberculosis* cultures grown in the absence of drugs (CT), or in the presence of M06 alone (1X MIC), of individual anti-tubercular drugs alone (2X MIC), or of their combinations: EMB ± M06 (**a**); MOX ± M06 (**b**). CFU are expressed as mean ± SD (2 ≤ *n* ≤ 3). **c−f**, Fraction of infected (**c**,**e**) and dead macrophages (**d**,**f**) at 6 days post infection treated with INH (2X MIC) ± M06 (1X MIC) (**c**,**d**) or with RIF (2X MIC) ± M06 (1X MIC) (**e**,**f**). UI (uninfected) and DMSO (untreated) macrophages are also reported. Error bars represent mean ± SD (*n* = 6): 1755 ≤ macrophages ≤ 2974 per condition (**c**,**d**); 2970 ≤ macrophages ≤ 4739 per condition (**e**,**f**). Significance by two-way ANOVA followed by Tukey’s multiple comparisons test: \**P* < 0.05; \*\**P* = 0.005; \*\*\**P* = 0.001; \*\*\*\**P* < 0.0001.

## References

1. Darby, E. M. et al. Molecular mechanisms of antibiotic resistance revisited. Nat Rev Microbiol (2022) doi:10.1038/s41579-022-00820-y.

2. Dewachter, L., Fauvart, M. & Michiels, J. Bacterial heterogeneity and antibiotic survival: Understanding and combatting persistence and heteroresistance. Molecular Cell 76, 255–267 (2019).

3. Verstraete, L., Van den Bergh, B., Verstraeten, N. & Michiels, J. Ecology and evolution of antibiotic persistence. Trends in Microbiology 30, 466–479 (2022).

4. Windels, E. M., Michiels, J. E., Van den Bergh, B., Fauvart, M. & Michiels, J. Antibiotics: Combatting tolerance to stop resistance. mBio 10, e02095–19 (2019).

5. Davis, K. M. For the greater (bacterial) good: Heterogeneous expression of energetically costly virulence factors. Infect Immun 88, e00911–19 (2020).

6. Urbaniec, J., Xu, Y., Hu, Y., Hingley-Wilson, S. & McFadden, J. Phenotypic heterogeneity in persisters: a novel ‘hunker’ theory of persistence. FEMS Microbiology Reviews 46, fuab042 (2022).

7. Theuretzbacher, U., Outterson, K., Engel, A. & Karlén, A. The global preclinical antibacterial pipeline. Nat Rev Microbiol 18, 275–285 (2020).

8. Defraine, V., Fauvart, M. & Michiels, J. Fighting bacterial persistence: Current and emerging anti-persister strategies and therapeutics. Drug Resistance Updates 38, 12–26 (2018).

9. Dartois, V. A. & Rubin, E. J. Anti-tuberculosis treatment strategies and drug development: challenges and priorities. Nat Rev Microbiol 20, 685–701 (2022).

10. Oh, S., Trifonov, L., Yadav, V. D., Barry, C. E. & Boshoff, H. I. Tuberculosis drug discovery: A decade of hit assessment for defined targets. Front. Cell. Infect. Microbiol. 11, 611304 (2021).

11. Edwards, B. D. & Field, S. K. The struggle to end a millennia-long pandemic: Novel candidate and repurposed drugs for the treatment of tuberculosis. Drugs 82, 1695–1715 (2022).

12. Gold, B. & Nathan, C. Targeting phenotypically tolerant *Mycobacterium tuberculosis*. Microbiol Spectr 5, 5.1.27 (2017).

13. Meylan, S., Andrews, I. W. & Collins, J. J. Targeting antibiotic tolerance, pathogen by pathogen. Cell 172, 1228–1238 (2018).

14. Huemer, M., Mairpady Shambat, S., Brugger, S. D. & Zinkernagel, A. S. Antibiotic resistance and persistence—Implications for human health and treatment perspectives. EMBO Reports 21, (2020).

15. Ernest, J. P. et al. Development of new tuberculosis drugs: Translation to regimen composition for drug-sensitive and multidrug-resistant tuberculosis. Annu. Rev. Pharmacol. Toxicol. 61, 495– 516 (2021).

16. Chung, E. S., Johnson, W. C. & Aldridge, B. B. Types and functions of heterogeneity in mycobacteria. Nat Rev Microbiol 20, 529–541 (2022).

17. Manina, G., Griego, A., Singh, L. K., McKinney, J. D. & Dhar, N. Preexisting variation in DNA damage response predicts the fate of single mycobacteria under stress. The EMBO Journal 38, (2019).

18. Ortseifen, V., Viefhues, M., Wobbe, L. & Grünberger, A. Microfluidics for biotechnology: Bridging gaps to foster microfluidic applications. Front. Bioeng. Biotechnol. 8, 589074 (2020).

19. Scheler, O., Postek, W. & Garstecki, P. Recent developments of microfluidics as a tool for biotechnology and microbiology. Current Opinion in Biotechnology 55, 60–67 (2019).

20. Mistretta, M., Gangneux, N. & Manina, G. Microfluidic dose–response platform to track the dynamics of drug response in single mycobacterial cells. Sci Rep 12, 19578 (2022).

21. Lima-Noronha, M.A. et al. Sending out an SOS - the bacterial DNA damage response. Genet Mol Biol 45, e20220107 (2022).

22. Ayotte, Y. et al. Fragment-based phenotypic lead discovery to identify new drug seeds that target infectious diseases. ACS Chem. Biol. 16, 2158–2163 (2021).

23. Coulibaly, S. et al. Phenanthrolinic analogs of quinolones show antibacterial activity against *M. tuberculosis*. European Journal of Medicinal Chemistry 207, 112821 (2020).

24. Blower, T. R., Williamson, B. H., Kerns, R. J. & Berger, J. M. Crystal structure and stability of gyrase–fluoroquinolone cleaved complexes from *Mycobacterium tuberculosis*. Proc. Natl. Acad. Sci. U.S.A. 113, 1706–1713 (2016).

25. Zhou, X., Ma, Z., Dong, D. & Wu, B. Arylamine N-acetyltransferases: a structural perspective: Understanding of NATs with their 3D structures. Br J Pharmacol 169, 748–760 (2013).

26. Payton, M., Auty, R., Delgoda, R., Everett, M. & Sim, E. Cloning and characterization of arylamine N-acetyltransferase genes from *Mycobacterium smegmatis* and *Mycobacterium tuberculosis*: Increased expression results in isoniazid resistance. J Bacteriol 181, 1343–1347 (1999).

27. Bhakta, S. et al. Arylamine N-acetyltransferase is required for synthesis of mycolic acids and complex lipids in *Mycobacterium bovis* BCG and represents a novel drug target. Journal of Experimental Medicine 199, 1191–1199 (2004).

28. Sim, E. et al. Arylamine N-acetyltransferases in mycobacteria. CDM 9, 510–519 (2008).

29. Palmer, A. C. & Kishony, R. Opposing effects of target overexpression reveal drug mechanisms. Nat Commun 5, 4296 (2014).

30. Abuhammad, A. et al. Structure of arylamine *N* -acetyltransferase from *Mycobacterium tuberculosis* determined by cross-seeding with the homologous protein from *M. marinum*: triumph over adversity. Acta Crystallogr D Biol Crystallogr 69, 1433–1446 (2013).

31. Feng, L., et al. The pentapeptide-repeat protein, MfpA, interacts with mycobacterial DNA gyrase as a DNA T-segment mimic. Proc. Natl. Acad. Sci. U.S.A. 118, e2016705118 (2021).

32. Manina, G., Dhar, N. & McKinney, J. D. Stress and host immunity amplify *Mycobacterium tuberculosis* phenotypic heterogeneity and induce nongrowing metabolically active forms. Cell Host & Microbe 17, 32–46 (2015).

33. Melin, J. & Quake, S. R. Microfluidic large-scale integration: The evolution of design rules for biological automation. Annu. Rev. Biophys. Biomol. Struct. 36, 213–231 (2007).

34. Scott, S. & Ali, Z. Fabrication methods for microfluidic devices: An overview. Micromachines 12, 319 (2021).

35. Nakagawa, S. & Cuthill, I. C. Effect size, confidence interval and statistical significance: a practical guide for biologists. Biological Reviews 82, 591–605 (2007).

36. Abbas-Aghababazadeh, F., Lu, P. & Fridley, B. L. Nonlinear mixed-effects models for modeling in vitro drug response data to determine problematic cancer cell lines. Sci Rep 9, 14421 (2019).

37. Cutler, K. J. et al. Omnipose: a high-precision morphology-independent solution for bacterial cell segmentation. Nat Methods 19, 1438–1448 (2022).

38. Lu, Y. et al. Screening for gene expression fluctuations reveals latency-promoting agents of HIV. Proc. Natl. Acad. Sci. U.S.A. 118, e2012191118 (2021).

39. Rego, E. H., Audette, R. E. & Rubin, E. J. Deletion of a mycobacterial divisome factor collapses single-cell phenotypic heterogeneity. Nature 546, 153–157 (2017).

40. Salaikumaran, M. R., Badiger, V. P. & Burra, V. L. S. P. 16S rRNA methyltransferases as novel drug targets against tuberculosis. Protein J 41, 97–130 (2022).

41. Laborde, J., Deraeve, C. & Bernardes-Génisson, V. Update of antitubercular prodrugs from a molecular perspective: Mechanisms of action, bioactivation pathways, and associated resistance. ChemMedChem 12, 1657–1676 (2017).

42. Kim, D.-W. et al. Identification of the enzyme responsible for *N*-acetylation of norfloxacin by *Microbacterium* sp. Strain 4N2-2. Appl Environ Microbiol 79, 314–321 (2013).

43. Abuhammad, A. et al. Piperidinols that show anti-tubercular activity as inhibitors of arylamine N-acetyltransferase: An essential enzyme for mycobacterial survival inside macrophages. PLoS ONE 7, e52790 (2012).

44. Sambandan, D. et al. Keto-mycolic acid-dependent pellicle formation confers tolerance to drug-sensitive *Mycobacterium tuberculosis*. mBio 4, e00222–13 (2013).

45. Madacki, J. et al. Impact of the epoxide hydrolase EphD on the metabolism of mycolic acids in mycobacteria. Journal of Biological Chemistry 293, 5172–5184 (2018).

46. Miller, C. et al. SOS response induction by β-Lactams and bacterial defense against antibiotic lethality. Science 305, (2004).

47. Dwyer, D. J., Collins, J. J. & Walker, G. C. Unraveling the physiological complexities of antibiotic lethality. Annu. Rev. Pharmacol. Toxicol. 55, 313–332 (2015).

48. Li, H. et al. Reactive oxygen species in pathogen clearance: The killing mechanisms, the adaption response, and the side effects. Front. Microbiol. 11, 622534 (2021).

49. Aubry, A., Pan, X.-S., Fisher, L. M., Jarlier, V. & Cambau, E. *Mycobacterium tuberculosis* DNA gyrase: Interaction with quinolones and correlation with antimycobacterial drug activity. Antimicrob Agents Chemother 48, 1281–1288 (2004).

50. Luan, G., Hong, Y., Drlica, K. & Zhao, X. Suppression of reactive oxygen species accumulation accounts for paradoxical bacterial survival at high quinolone concentration. Antimicrob Agents Chemother 62, e01622–17 (2018).

51. Nepali, K., Lee, H.Y. & Liou, J.P. Nitro-group-containing drugs. J Med Chem 62, 2851–2893 (2019).

52. Atmane, N., Dairou, J., Paul, A., Dupret, J.-M. & Rodrigues-Lima, F. Redox regulation of the human xenobiotic metabolizing enzyme arylamine N-acetyltransferase 1 (NAT1). Journal of Biological Chemistry 278, 35086–35092 (2003).

53. VanDrisse, C. M. & Escalante-Semerena, J. C. Protein acetylation in bacteria. Annu. Rev. Microbiol. 73, 111–132 (2019).

54. Bienvenut, W. V. et al. Dual lysine and N-terminal acetyltransferases reveal the complexity underpinning protein acetylation. Molecular Systems Biology 16, (2020).

55. Kim, J.-E., Choi, J.-S., Kim, J.-S., Cho, Y.-H. & Roe, J.-H. Lysine acetylation of the housekeeping sigma factor enhances the activity of the RNA polymerase holoenzyme. Nucleic Acids Research 48, 2401–2411 (2020).

