## Supplemental Information for "Dynamic microfluidic single-cell screening identifies pheno-tuning compounds to potentiate tuberculosis therapy"

**This PDF file includes:**

**Supplementary Figures 1 to 3**
**Supplementary Table legends 1 to 6**
**Supplementary Table 7**
**Supplementary Video legends 1 and 2**
**Supplementary Methods**
**Supplementary References 56 to 72**

### Supplementary Figures

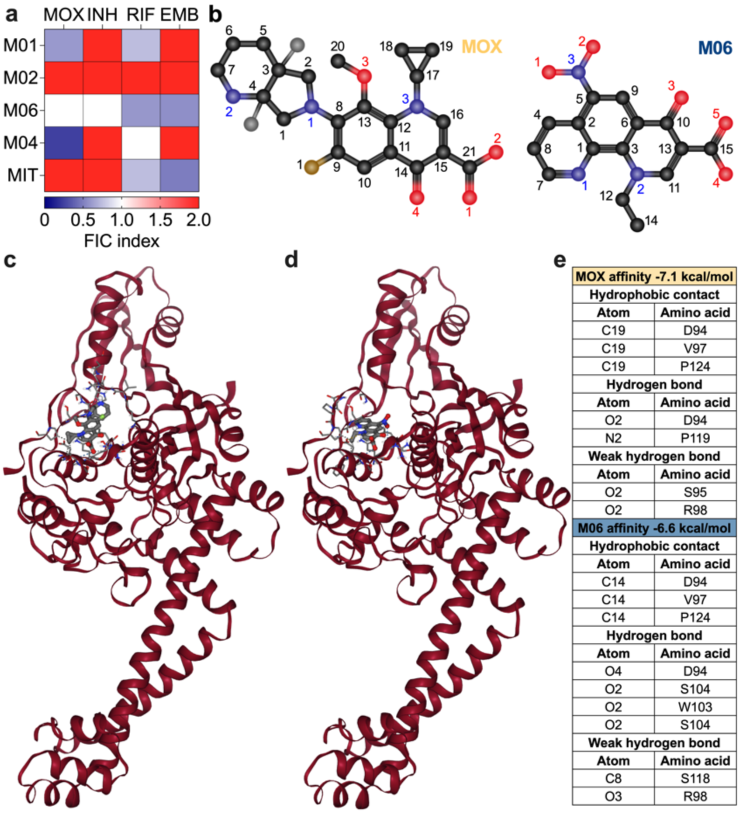


**Supplementary Figure 1. In-vitro and in-silico analyses of PTC activity against *M. tuberculosis*.** **a**, Heatmap displaying fractional inhibitory concentration (FIC) indices calculated for combinations of PTC and MIT with anti-tubercular drugs: indifference (FIC > 1); additivity (1 < FIC < 0.5), synergy (FIC ≤ 0.5). Experiments were repeated at least twice independently. **b**, CPK-colored chemical structures of MOX and M06. Molecular topology is also indicated. **c**,**d**,**e**, In silico docking poses of MOX (**c**) and M06 (**d**) in *M. tuberculosis* DNA gyrase, and table of predicted molecular interactions (**e**).

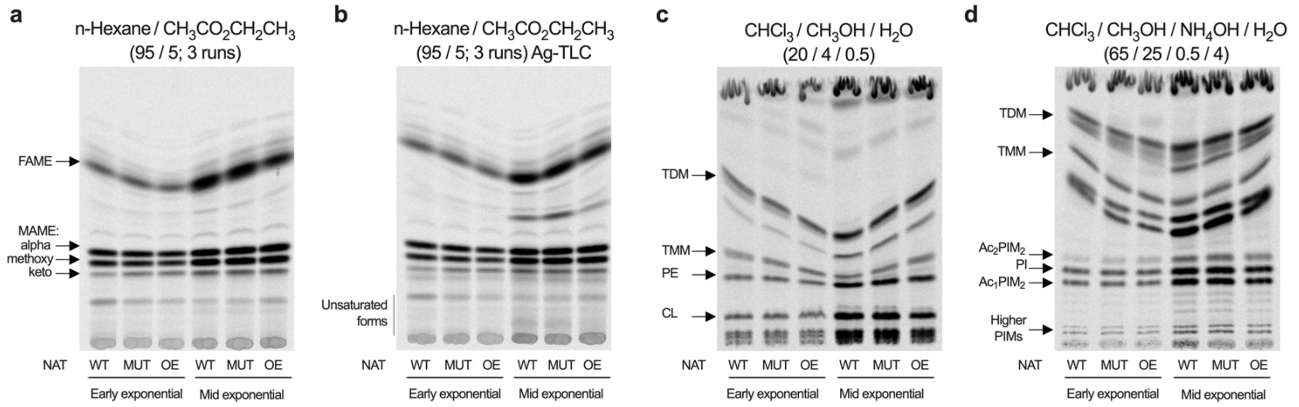


**Supplementary Figure 2. Analysis of lipids in *M. tuberculosis* expressing different NAT variants. a−d**, Representative TLC of fatty and mycolic acids and lipids extracted from *M. tuberculosis* WT; S10 mutant (MUT); and NAT overexpressing strain (OE) in early and mid-exponential phase, after 24-hour incorporation of $^{14}$C-acetate (1 µCi/mL). Conditions for separation of different lipid populations are shown above the TLC autoradiographs. Fatty acid methyl esters (FAME) and mycolic acid methyl esters (MAME) (**a**). Unsaturated forms of MAME (**b**). Trehalose dimycolate (TDM); trehalose monomycolate (TMM); phosphatidylethanolamine (PE); cardiolipin (CL) (**c**). TDM; TMM; phosphatidylinositol (PI); phosphatidylinositol mannosides (PIM); acylated forms of PIM ($Ac_{2/1}PIM_2$) (**d**).

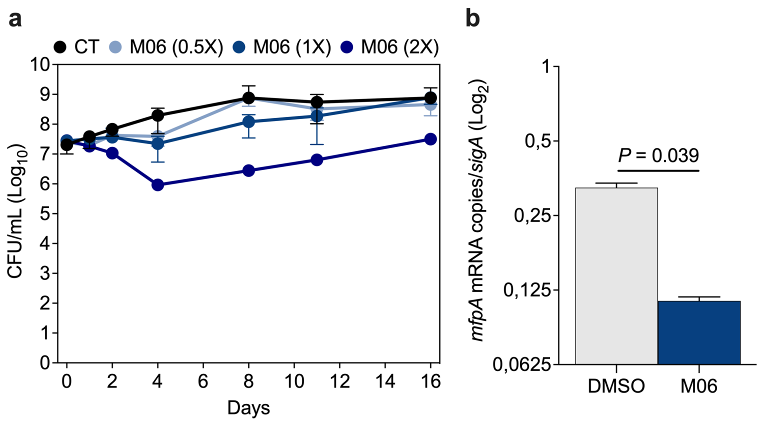


**Supplementary Figure 3. Effect of M06 alone in vitro. a**, M06 activity against exponential-phase *M. tuberculosis* cultures grown in the absence of drugs (CT), or in the presence of M06 alone at 0.5X, 1X, or 2X the MIC (1.25 µg/mL). Error bars represent mean ± SD ($n$ = 3). **b**, qRT-PCR analysis of *M. tuberculosis rv3361c* (*mfpA*) after DMSO or M06 treatment (10X MIC) for 24 hours. Transcripts are normalized to total RNA and *sigA* copies and are expressed as $\log_2$ fold change before and after treatment. Error bars represent mean ± SD ($n$ = 2). Significance by paired $t$-test.

### Supplementary Tables

**Supplementary Table 1. PTC candidates used for the screening.** List of fragment compounds and phenanthroline derivatives used in this study (Fig. 2). Structure, name, molecular weight, MIC in mycobacteria, and IC50 in Vero cells are reported.

**Supplementary Table 2. Screening results, extracted parameters and microcolony indices.** Output datasheets derived from the analysis of 15 µDeSCRiPTor experiments (Fig. 2). Each experiment is identified by an alphanumeric code and three files per experiment are present, reporting different parameters over time: microcolony area in pixels (area); $\log_{10}$ transformed RecA-GFP mean fluorescence (mean); and $\log_{10}$ transformed fluorescence variance (var) in arbitrary units. Column B of each datasheet reports time relative to the onset of drug injection, in hours. Column C indicates the absence or presence of treatment. The following columns report the parameter for different colonies, identified in the first row by condition and number. The following datasheet, named cent_results, reports an assembly of all screening results listed in previous datasheets as follows: experiment ID; condition; exposure stage; average of each index across the four last time points of the PRE stage and across the four last time points of the DUR stage for each microcolony. The last two datasheets, named effect_sizes_mean or var, contain the outcome of effect size estimations.

**Supplementary Table 3. Differential analysis of *M. tuberculosis* transcriptome upon PTC exposure.** Raw and normalized counts per gene in each condition tested and in the control condition DMSO ($n$ = 3). Fold changes and relative statistics, and Benjamini-Hochberg corrected $P$ values (padj) are also reported for each comparison. Two datasheets are provided for each PTC hit, color-coded as in Fig. 3. The first datasheet includes upregulated genes (UP) and the second one includes downregulated genes (DN).

**Supplementary Table 4. Functional and GO enrichment analyses of DEGs upon PTC exposure.** Two datasheets are included for each condition: functional analysis (FA) and gene ontology enrichment (GO) (Fig. 3 and Extended Data Fig. 3). Each FA datasheet reports merged differential analysis of upregulated (black font, UP) and downregulated (white font, DN) genes, sorted by functional category using Mycobrowser's color coding (https://mycobrowser.epfl.ch). Each GO datasheet reports GO biological processes identified by PANTHER overrepresentation test of significant DEGs against all genes in *M. tuberculosis* H37Rv database[70,71]. Significance was assessed by Fisher's exact test followed by Benjamini-Hochberg procedure, filtering valid results for FDR < 0.05. Results are sorted by fold enrichment of the most specific categories (highlighted), with their parent terms indented below. Datasheet names include the type of analysis, the condition, and the direction of DEGs.

**Supplementary Table 5. List and features of *gyrA* point mutants and multi-drug resistant strains.** The table shows the source, identifier, name, susceptibility profile and associated mutations of strains shown in Fig. 3e.

**Supplementary Table 6. Single nucleotide polymorphisms (SNP) of PTC-resistant mutants.** Results of variant calling analysis relative to four spontaneous mutants isolated on M02 or M06. Features and effects of each SNP are reported, compared to the parent strain. A summary of these results is shown in Fig. 4a.

**Supplementary Table 7. Strains, plasmids, and primers used in this study.**

| REAGENT | SOURCE | IDENTIFIER |
|---|---|---|
| **Bacterial strains and Cell lines** | | |
| *Escherichia coli* TOP10 | ThermoFisher | #C404010 |
| *Mycobacterium smegmatis* mc$^2$155 | Lab collection | ATCC 700084 |
| *Mycobacterium tuberculosis* Erdman | Lab collection | ATCC 35801 |
| *Mycobacterium tuberculosis* H37Rv | Lab collection | ATCC 27294 |
| ATCC 700084 transcriptional reporter of *recA* (*MSMEG_2723*), with destabilized GFP under the control of native *recA* promoter | Lab collection[17] | GMS2 |
| GMS2 constitutively expressing wild type mCherry | This paper | GMS2_pGM218 |
| ATCC 27294/pGM321 constitutively expressing *nat* (*rv3566c*) | This paper | GMT35 |
| ATCC 27294/pGM315; *nat* (*rv3566c*) CRISPRi/dCas9 | This paper | GMT36 |
| ATCC 35801 transcriptional reporter of *rrs*, with destabilized GFP under the control of native rRNA promoter, also constitutively expressing DsRed2 | Lab collection[32] | GMT17 |
| ATCC 27294 carrying gyrase point mutation (A90V) | CIP111339 | PM_1 |
| ATCC 27294 carrying gyrase point mutation (S91P) | CIP111343 | PM_2 |
| ATCC 27294 carrying gyrase point mutation (G88C) | CIP111344 | PM_3 |
| *M. tuberculosis* multi-drug resistant clinical isolate | CIP111418 | MDR_4 |
| *M. tuberculosis* multi-drug resistant clinical isolate | CIP111386 | MDR_5 |
| *M. tuberculosis* multi-drug resistant clinical isolate | CIP111390 | MDR_6 |
| *M. tuberculosis* multi-drug resistant clinical isolate | CIP111392 | MDR_7 |
| *M. tuberculosis* multi-drug resistant clinical isolate | CIP111393 | MDR_8 |
| RAW 264.7 macrophages | ATCC | TIB-71 |
| Vero cells | ATCC | CCL-81 |

| | | |
|---|---|---|
| **Plasmids** | | |
| pCR2.1-TOPO TA cloning plasmid, Amp$^{R}$, Km$^{R}$ | Invitrogen | pCR2.1-TOPO |
| pND200 expressing mCherry from UV15 strong promoter, Hyg$^{R}$ | Lab collection | pGM218 |
| pLJR965, tetracycline-inducible dCas9 *attB*-integrative vector for *M. tuberculosis* gene silencing, Km$^{R}$ | Addgene[72] | #115163 |
| pLJR965 modified with Hyg$^{R}$ cassette | Lab collection | pGM309 |
| pGM309 carrying sgRNA for *nat* (*rv3566c*) silencing, Hyg$^{R}$ | This paper | pGM315 |
| pND200 expressing *nat* (*rv3566c*) from UV15 strong promoter, Hyg$^{R}$ | This paper | pGM321 |
| **Primers** | | |
| siRNA targeting sequence, sgRNA_*nat*_F (*rv3566c*)<br>5'-GGGAGGACTGTGCACGGTCACCAGATCCT-3' | This paper,<br>IDT | # 230084119 |
| siRNA targeting sequence, sgRNA_*nat*_R (*rv3566c*)<br>5'-AAACAGGATCTGGTGACCGTGCACAGTCC-3' | This paper,<br>IDT | # 230084120 |
| CT primer silencing, pCRISPRi_Seq | Lab collection<br>Eurofins | # 26983185 |
| CT primer silencing, pLJR_PCR – *for* | Lab collection<br>Eurofins | # 27008276 |
| Cloning, SD-*rv3566c*, NAT_oe – *for*<br>5'-AC**TTAATTAA**GAAGGAGATATACAT<br>ATGGCACTGGATCTGACCGCG-3' | This paper,<br>IDT | # 232873379 |
| Cloning, SD-*rv3566c*, NAT_oe – *rev*<br>5'-AT**GTTAAC**TTACGGCGCATCGGCTCCTGG-3' | This paper,<br>IDT | # 232873380 |
| qPCR*, sigA (rv2703) – for*<br>5'-ACGACGAAGACCACGAAGAC-3' | This paper,<br>Eurofins | # 27779941 |
| qPCR*, sigA (rv2703) – rev*<br>5'-CTTCATCCCAGACGAAATCACC-3' | This paper,<br>Eurofins | # 27779942 |
| qPCR, *nat* (*rv3566c*) – *for*<br>5'-GGCTGATGGGTTATGTGCTG-3' | This paper,<br>Eurofins | # 32757720 |
| qPCR*, nat* (*rv3566c) – rev*<br>5'-ATCCGACGTCGACGAGATAG-3' | This paper,<br>Eurofins | # 32757721 |
| qPCR*, ephA* (*rv3617*) – *for*<br>5'-GGCTTTATCGATCGGCTTCC-3' | This paper,<br>Eurofins | # 33124397 |
| qPCR*, ephA* (*rv3617*) – *rev*<br>5'-GTGAACTCGCCGATGTAGTG-3' | This paper,<br>Eurofins | # 33124398 |

| | | |
|---|---|---|
| qPCR, *ephD* (*rv2214c*) – *for* 5’-TACATGGCCTTGTTCTCGGT-3’ | This paper, Eurofins | # 33124399 |
| qPCR, *ephD* (*rv2214c*) – *rev* 5’-AGCGTCTCCGAGTGATGAAT-3’ | This paper, Eurofins | # 33124401 |
| qPCR, *sodA* (*rv3846*) – *for* 5’-CAGCGATCTTGCTGAACGAA-3’ | This paper, Eurofins | # 33124401 |
| qPCR, *sodA* (*rv3846*) – *rev* 5’-CACGGAACTTGTCGAACGAA-3’ | This paper, Eurofins | # 33124402 |
| qPCR, *sodC* (*rv0432*) – *for* 5’-CCTTCACCATGGACGACCT-3’ | This paper, Eurofins | # 33124403 |
| qPCR, *sodC* (*rv0432) – rev* 5’-GTCCCATTGACCTGGACGTA-3’ | This paper, Eurofins | # 33124404 |
| qPCR, *trxB1*(*rv1471*) – *for* 5’-GACTACCCGAGACCTCACTG-3’ | This paper, Eurofins | # 33124405 |
| qPCR, *trxB1* (*rv1471*) – *rev* 5’-ACCAGGAGGCCCAATAATCG-3’ | This paper, Eurofins | # 33124406 |
| qPCR, *ahpE* (*rv2238*) – *for* 5’-CGACCAGAATCAGCAGCTTG-3’ | This paper, Eurofins | # 33124407 |
| qPCR, *ahpE* (*rv2238*) – *rev* 5’-CAACAGCACGTTCTTTGCAC-3’ | This paper, Eurofins | # 33124408 |
| qPCR, *katG* (*rv1908c) – for* 5’-AAGGCCTGGTACAAGCTGAT-3’ | This paper, Eurofins | # 33124409 |
| qPCR, *katG* (*rv1908c*) – *rev* 5’-GGCTCTTAAGGCTGGCAATC-3’ | This paper, Eurofins | # 33124410 |
| qPCR, *recA (rv2737c) – for* 5’-CACGACGACCTCTGAACAAC-3’ | This paper, Eurofins | # 33124413 |
| qPCR, *recA (rv2737c) – rev* 5’-CGTAATCTCGAACGGTGCTC-3’ | This paper, Eurofins | # 33124414 |
| qPCR, *mfpA (rv3361c) – for* 5’-CGTTGGACGACGTGGATTTC-3’ | This paper, Eurofins | # 33124415 |
| qPCR, *mfpA (rv3361c) – rev* 5’-TCAAGTTGAGACCACGCAGA-3’ | This paper, Eurofins | # 33124416 |

**Supplementary Videos**

**Supplementary Video 1. Time-lapse microscopy of wild-type *M. smegmatis* growing in the 32-condition platform.** Representative video of exponential-phase bacteria growing inside a random microchamber of the 32-condition platform and fed by constant flow of fresh 7H9 medium. Images were recorded at 20-minute intervals (10 fps). Time is shown in hours and minutes. Scale bar, 5 µm.

**Supplementary Video 2**. **Combined time-lapse microscopy of *M. smegmatis* RecA-GFP_mCherry$_{cyt}$ reporter during the µDeSCRiPTor.** Representative movies of exponentially growing bacteria seeded into the 32-condition platform during the screening. Bacteria were first grown in fresh 7H9 medium between 4 to 6 hours. Next, bacteria were treated for 6 hours with subinhibitory concentrations of control compounds (DMSO, MIT, MOX and INH); PTC hits (M01, M02, M04 and M06); or two PTC (B02 and F05) causing a significant but opposite effect to that of PTC hits. Finally, fresh 7H9 medium was perfused everywhere for 6 hours. Images were recorded every 30 minutes (10 fps). Phase contrast (red) and RecA-GFP (cyan) channels are merged. Time is shown in hours and minutes, and conditions are indicated. Scale bars, 5 µm.

### Supplementary Methods

Three phenanthroline derivatives (**M01 – MR34503; M05 – MR34504; M06 – MR34509**) were formerly published[23], whereas four of them were synthesized in this work. All chemical reagents and solvents were purchased from commercial sources and used without further purification. Melting points were determined on a Kofler melting point apparatus. $^{1}H$ and $^{13}C$ NMR spectra were recorded on a BRUKER AVANCE III 400 MHz with chemical shifts expressed in parts per million (in chloroform-*d*) downfield from TMS as an internal standard and coupling in Hertz. IR spectra were recorded on a PerkinElmer BX FT-IR apparatus using KBr pellets. High resolution mass spectra (HRMS) were obtained by electrospray on a BrukermaXis. The purities of all tested compounds were analyzed by LC–MS, with the purity all being higher than 95%. Analyses were performed with a Waters Alliance 2695 as separating module (column XBridge C18 2.5 mM/4.6 x 50 mM) using the following gradients: A (95%)/B (5%) to A (5%)/B (95%) in 4.00 min. This ratio was hold during 1.50 min before return to initial conditions in 0.50 min. Initial conditions were then maintained for 2.00 min (A ¼ $H_2O$, B ¼ $CH_3CN$; each containing HCOOH: 0.1%). MS were obtained on a SQ detector by positive ESI.

**3-(dimethylamino)propyl 1-methyl-6-nitro-4-oxo-1,4-dihydro-1,10-phenanthroline-3-carboxylate (M02 – MR36009):** From 1-methyl-6-nitro-4-oxo-1,10-phenanthroline-3-carboxylic acid(1 eq, 0.33 mmol, 100 mg), dry THF (4 mL), oxalyl chloride (2.5 eq, 0.82 mmol, 69 µL) and DMF (2 drops) according to the general procedure A. The mixture was stirred for 3h. After concentration under pressure, DCM (4 mL), triethylamine (1.3 eq, 0.43 mmol, 60 µL) and 3-(dimethylamino)propan-1-ol (1.3 eq, 0.43 mmol, 51 µL) were stirred for 30 min. The product was washed with sodium carbonate satured water. The organic layer was dried over $MgSO_4$. Removal of the solvant under reduced pressure afforded the crude product, which was purified by chromatography on silica gel column (DCM/MeOH/$NH_3$, gradient 100:0:0 to 80:20:0.2) to give the compound (24%).$^{1}H$ NMR($CDCl_3$, 400 MHz) δ 9.26 (s, 1H), 9.06 (s, 1H), 9.04 (dd, *J* = 3 Hz, 1H), 8.54 (s, 1H), 7.82 – 7.73 (m, 1H), 4.66 (s, 3H), 4.40 (t, *J* = 6.7 Hz, 2H), 2.51 (t, *J* = 7.3 Hz, 2H), 2.29 (s, 6H), 1.99 (p, J= 6.8 Hz, 2H).13C NMR ($CDCl_3$, 100 MHz) δ 172.3, 164.7, 153.3, 148.5, 142.1, 141.7, 141.3, 132.5, 127.6, 124.7, 123.7, 123.4, 114.4, 63.8, 56.3, 50.6, 45.5, 26.9. MS m/z $[M+H]^+$ 384.87. IR (neat, cm-1) 3063, 2953, 2816, 2766, 2242, 1696, 1639, 1599, 1520, 1501, 1341, 1204, 1089. Mp156.8°C. HRMS m/z $[M+H]^+$ calc 385.1512 exp 385.1513. Mp 156.8°C

**2-bromoethyl 1-methyl-6-nitro-4-oxo-1,4-dihydro-1,10-phenanthroline-3-carboxylate (M03 – MR36013):** From 1-methyl-6-nitro-4-oxo-1,10-phenanthroline-3-carboxylic acid (1 eq, 0.23 mmol, 69 mg), dry THF (3 mL), oxalyl chloride (2 eq, 0.46 mmol, 39 µL) and DMF (2 drops) according to the general procedure A. After concentration under pressure, DCM (3 mL), triethylamine (1.3 eq, 0.30 mmol, 42 µL) and 2-bromoethanol (1.3 eq, 0.30 mmol, 21 µL) were stirred for 1 h. DCM was then added and the precipitate was filtered under vacuum. Removal the solvent of filtrate under reduced pressure afforded the product, which was purified by chromatography on silica gel column (DCM/AcOEt, gradient 100:0 to 50:50) to give a yellow bright solid (20%). $^{1}$H NMR ($CDCl_3$, 400 MHz) δ 9.37 (s, 1H), 9.15 –9.06 (m, 2H), 8.60 (s, 1H), 7.80 (dd, *J* = 8.8, 4.2 Hz, 1H), 4.76 –4.64 (m, 5H), 3.69 (t, *J* = 6.3 Hz, 2H).$^{13}$C NMR ($CDCl_3$, 100 MHz) δ 172.4, 164.0, 153.5, 148.6, 142.4, 141.8, 141.4, 132.7, 127.9, 124.8, 123.9, 123.5, 113.8, 64.5, 50.8, 28.8. MS m/z $[M+H]^+$ 405.68 - 407.68. IR (neat, cm-1) 3067, 2931, 1734, 1690, 1640, 1622, 1518, 1478, 1370, 1319, 1255, 1198, 1118, 1085. Mp 234.1°C.

***N*, *N*, 1-trimethyl 6-nitro-4-oxo-1,4-dihydro-1,10-phenanthroline-3-carboxylamide (M04 – MR36018):** From 1-methyl-6-nitro-4-oxo-1,10-phenanthroline-3-carboxylic acid (1 eq, 0.23 mmol, 69 mg), dry THF (5 mL), oxalyl chloride (2 eq, 0.46 mmol, 39 µL) and DMF (2 drops) according to the general procedure A. After concentration under pressure, DCM (5 mL), triethylamine (1.3 eq, 0.3 mmol, 42 µL) and dimethylamine (1.3 eq, 0.3 mmol, 15 µL) were stirred for 1 h. DCM was then added and the organic phase was washed once with $NaHCO_3$ sat, dried on $MgSO_4$ and concentrated. The crude product was purified by chromatography on desactivated silica gel column (DCM/MeOH, gradient 100:0 to 80:20) to give a yellow solid (50%). $^{1}$H NMR ($CDCl_3$, 400 MHz) δ 9.37 (s, 1H), 9.14 (dd, *J* = 8.8, 1.7 Hz, 1H), 9.07 (dd, *J* = 4.2, 1.7 Hz, 1H), 8.09 (s, 1H), 7.80 (dd, *J* = 8.8, 4.2 Hz, 1H), 4.65 (s, 3H), 3.16 (s, 3H), 3.07 (s, 3H).$^{13}$C NMR ($CDCl_3$,100 MHz) δ 171.8, 166.2, 149.6, 148.3, 141.9, 141.8, 141.5, 132.7, 125.9, 124.6, 123.8, 123.8, 123.0, 50.2, 38.7, 35.8. MS m/z $[M+H]^+$ 326.91. IR (neat, cm-1) 2963, 2922, 2852, 1719, 1636, 1501, 1384, 1261, 1078.

**1-(dimethylamino)prop-2-yl 1-methyl-6-nitro-4-oxo-1,4-dihydro-1,10-phenanthroline-3-carboxylate (M07 – MR36017):** From 1-methyl-6-nitro-4-oxo-1,10-phenanthroline-3-carboxylic acid (1 eq, 0.27 mmol, 80 mg), dry THF (5 mL), oxalyl chloride (2 eq, 0.54 mmol, 45 µL) and DMF (2 drops) according to the general procedure A. After concentration under pressure, DCM (5 mL), triethylamine (1.3 eq, 0.35 mmol, 49 µL) and 1-dimethylaminopropan-2-ol (1.3 eq, 0.36 mmol, 43 µL) were stirred for 1 h. DCM was then added and the organic phase was washed once with $NaHCO_3$ sat, dried on $MgSO_4$ and concentrated. The crude product was purified by chromatography on desactivated silica gel column (DCM/MeOH, gradient 100:0 to 80:20) to give a yellow solid (44%). $^{1}$H NMR ($CDCl_3$, 400 MHz) δ 9.35 (s, 1H), 9.10 (dd, *J* = 8.8, 1.7 Hz, 1H), 9.06 (dd, *J* = 4.2, 1.7 Hz, 1H), 8.55 (s, 1H), 7.78 (dd, *J* = 8.8, 4.1 Hz, 1H), 5.41 – 5.28 (m,1H), 4.66 (s, 3H), 2.73 (dd, J = 12.9, 7.3 Hz, 2H), 2.42 (dd, J = 12.9, 5.1 Hz, 1H), 2.32 (s, 6H), 1.40 (d, J = 6.3 Hz, 3H).$^{13}$C NMR ($CDCl_3$,100 MHz) δ 172.5, 164.2, 153.2, 148.5, 142.1, 141.8, 141.3, 132.7, 127.8, 124.7, 123.8, 123.6, 114.9, 69.7, 64.4, 50.5, 46.3, 46.2, 18.8. MS m/z $[M+H]^+$ 384.94. IR (neat, cm-1) 1712, 1630, 1597, 1501, 1335, 1264, 1200, 1082. HRMS m/z $[M+H]^+$ calc 385.1512 exp 385.1515. Mp 119.7°C

**Supplementary References**


56. Schneider, C.A., Rasband, W.S., & Eliceiri, K.W. NIH Image to ImageJ: 25 years of image analysis. *Nat Methods* **9**, 671–675 (2012).
57. Murail, S., de Vries, S. J., Rey, J., Moroy, G. & Tufféry, P. SeamDock: An interactive and collaborative online docking resource to assist small compound molecular docking. *Front. Mol. Biosci.* **8**, 716466 (2021).
58. Cokelaer, T., Desvillechabrol, D., Legendre, R. & Cardon, M. 'Sequana': a Set of Snakemake NGS pipelines. *JOSS* **2**, 352 (2017).
59. Köster, J. & Rahmann, S. Snakemake—a scalable bioinformatics workflow engine. *Bioinformatics* **28**, 2520–2522 (2012).
60. Li, H. & Durbin, R. Fast and accurate short read alignment with Burrows–Wheeler transform. *Bioinformatics* **25**, 1754–1760 (2009).
61. Tarasov, A., Vilella, A. J., Cuppen, E., Nijman, I. J. & Prins, P. Sambamba: fast processing of NGS alignment formats. *Bioinformatics* **31**, 2032–2034 (2015).
62. Sandmann, S. *et al.* appreci8: a pipeline for precise variant calling integrating 8 tools. *Bioinformatics* **34**, 4205–4212 (2018).
63. Cingolani, P. *et al.* A program for annotating and predicting the effects of single nucleotide polymorphisms, SnpEff: SNPs in the genome of Drosophila melanogaster strain $w^{1118}$; iso-2; iso-3. *Fly* **6**, 80–92 (2012).
64. Ewels, P., Magnusson, M., Lundin, S. & Käller, M. MultiQC: summarize analysis results for multiple tools and samples in a single report. *Bioinformatics* **32**, 3047–3048 (2016).
65. Martin, M. Cutadapt removes adapter sequences from high-throughput sequencing reads. *EMBnet.journal*, **17**, 10–12 (2011).
66. Langmead, B. & Salzberg, S.L. Fast gapped-read alignment with Bowtie 2. *Nat Methods* **9**, 357–359 (2012).
67. Liao, Y., Smyth, G. & Shi, W. FeatureCounts: An efficient general purpose program for assigning sequence reads to genomic features. *Bioinformatics* **30**, 923–930 (2014).
68. Love, M.I., Huber, W. & Anders, S. Moderated estimation of fold change and dispersion for RNA-seq data with DESeq2. *Genome Biol* **15**, 550 (2014).
69. Varet, H., Brillet-Guéguen, L., Coppée, J.Y. & Dillies, M.A. SARTools: A DESeq2- and EdgeR-Based R pipeline for comprehensive differential analysis of RNA-Seq data. *PLoS One* **11**, e0157022 (2016).
70. The Gene Ontology Consortium *et al.* The Gene Ontology resource: enriching a GOld mine. *Nucleic Acids Research* **49**, D325–D334 (2021).
71. Mi, H., Muruganujan, A., Ebert, D., Huang, X. & Thomas, D. PANTHER version 14: more genomes, a new PANTHER GO-slim and improvements in enrichment analysis tools. *Nucleic Acids Research* **47**, D419–D426 (2019).
72. Rock, J. M. *et al.* Programmable transcriptional repression in mycobacteria using an orthogonal CRISPR interference platform. *Nat Microbiol* **2**, 16274 (2017).
